# Locomotion optimizes sensory representations through a computational principle shared by rodents and primates

**DOI:** 10.1101/2025.06.29.662230

**Authors:** Jonathan M. Gant, Wiktor F. Młynarski

## Abstract

Behavior modulates the activity of sensory systems in multiple ways: from gain changes in individual neurons to changing interactions in neural populations. These effects are not universal; while movement has a strong influence on sensory coding in rodents, its impact on primates is less prominent. The diversity of effects that locomotion exerts on sensory neurons, as well as disparities between species, raises questions about the existence of universal principles that may underlie sensation during behavior. We propose that sensory systems are internally modulated to match systematic changes in stimulus statistics caused by locomotion, to facilitate an accurate and efficient sensory code. We find that model neurons, adapted to stimuli recorded during movement in natural environments, predict and reproduce a broad spectrum of experimental observations in rodents and primates. This simple principle of maintaining coding efficiency across behavioral states reconciles the diversity of ways in which locomotion modulates visual coding in different animal species.

## INTRODUCTION

Behavior exerts a substantial modulatory influence on brain regions traditionally regarded as purely sensory (*1*). Locomotion, in particular, has been shown to alter the gain of visual responses across multiple species, including fruit flies (*2, 3*) and mice (*4, 5*). In rodents, movement induces more complex changes in the visual system, modulating not only the response strength but also the temporal dynamics of neural activity (*6*), spatial integration of stimuli (*7* ), and the correlation structure of neural populations (*8*). These findings challenge the classical notion of sensory processing as a behaviorally invariant computation, highlighting instead the dynamic interplay between sensory encoding and behavioral state.

Recent experimental findings reveal an even more complex relationship between sensory coding and movement: neurons in the primary visual cortex (V1) of primates are modulated by running in a different way than those of mice (*9*). These disparities raise intriguing questions regarding whether strategies of visual coding during movement are universal or species-specific, and whether they could be understood with the same principles.

In this work, we take a normative perspective and consider how locomotion should modulate sensory neurons such that stimuli are encoded accurately and efficiently in different behavioral states. Our approach is grounded in the efficient coding hypothesis, an established theoretical framework that proposes design objectives for sensory coding (*10*–*12*). The hypothesis postulates that sensory systems adapt to the statistical structure of stimuli in order to maximize the amount of information transmitted, minimize encoding errors, and satisfy metabolic constraints. According to this theoretical perspective, at the onset of movement, the brain should adjust sensory neurons to systematic changes in input statistics caused by behavior. This strategy would ensure that stimuli are encoded efficiently and accurately in different behavioral states.

To understand how sensory input changes with locomotion in the natural environment, we recorded videos of natural scenes with a stationary and moving camera. We processed these videos with filters designed to mimic the receptive fields in V1 of mice and primates. We then developed models of sensory coding that are optimally modulated to match different statistical features of natural stimuli recorded during movement and rest. Optimized model neurons capture a range of phenomena observed in the mouse visual system: gain (*4*) and tuning curve modulation (*8*), changes in encoding accuracy (*13*), as well as population-level interactions that modulate spatial integration (*14*). By analyzing the temporal structure of natural stimuli during movement, we formed predictions about the dynamics of neural activity during locomotion and confirmed them by reanalyzing the experimental data of (*6*). To confirm that the observed changes in stimulus statistics are a part of the animal’s experience during natural behavior, we reanalyzed visual input data from the perspective of freely moving mice, made available in (*15*). Going beyond the visual system of rodents, we found that neurons with high spatial frequency receptive fields, such as those in the foveal area of primate V1, do not benefit from modulation because their statistics do not change sufficiently strongly with the animal’s movement. Our findings indicate that the diverse and seemingly disparate effects of locomotion on sensory coding share a common general principle: maintain the accuracy and efficiency of information transmission across different behavioral states.

## RESULTS

The proposed principle of sensory coding during locomotion is based on two observations (Fig. 1a). First, locomotion substantially impacts the sensory experience of the organism by changing the statistics of sensory input compared to the stationary state (Fig. 1a, thick black arrow). Second, if this impact is systematic and predictable, the brain can preemptively adjust (i.e. modulate) sensory systems at the onset of movement such that they are prepared to efficiently process stimuli whose statistics are impacted by locomotion (Fig. 1a, thick orange arrow).

**Figure 1:**
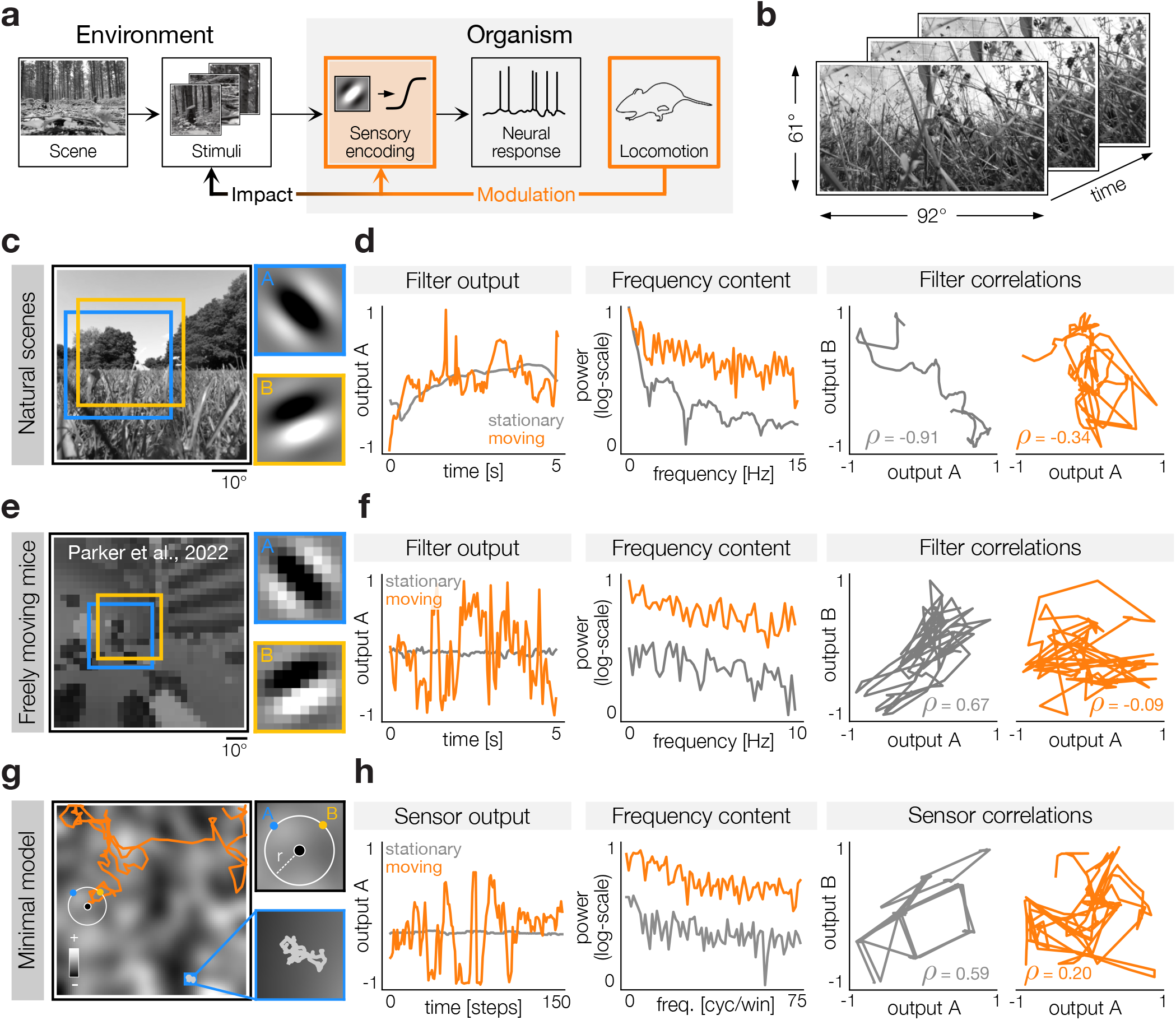
Internal modulation adjusts sensory neurons to changes in stimulus statistics caused by locomotion. **a)** Schematic of the proposed modulation principle. Sensory encoding maps stimuli (e.g. images) on neural activity. Locomotion impacts the statistics of stimuli and it should modulate sensory systems to maintain efficiency of the sensory code. **b)** Recordings of natural scenes during movement and in a stationary condition. **c)** A frame from an example natural video (left, cropped for visualization) processed with two Gabor filters (right). The size and position of the filters are marked with yellow and blue frames (left). **d)** Changes in filter outputs during stationary periods (gray) and camera movement (orange). Left: temporal trajectories of filter outputs differ in dynamic range. Middle: locomotion impacts the frequency spectrum of filter outputs. Right: locomotion reduces the magnitude of correlation between filter outputs. **e)** Visual input from the perspective of a freely moving mouse (reanalyzed data from (*15*)). A frame from an example video. The videos were processed with Gabor filters (right) with the same parameters as in (c). **f)** In freely moving mice, the impact of locomotion on stimulus statistics is similar to the effect in natural videos. Left, middle and right panels correspond to the same panels in (d). **g)** A minimal model of stimulus changes due to locomotion. An agent with two sensors (left, top-right; black, blue and yellow dots respectively) moves in a simulated two-dimensional environment (left, orange line) or fluctuates around the same location, which corresponds to a stationary state (left and bottom right, gray trajectory, blue area). **h)** The minimal model recapitulates changes in sensory inputs due to locomotion in natural scenes with complex statistics. Left, middle and right panels correspond to the same panels in (d).

A key assumption is that natural selection shaped sensory systems over evolutionary timescales to exploit the statistics of natural scenes which change with locomotion in a predictable way. The resulting modulation of sensory systems should therefore persist even in experimental settings that are not entirely natural. This principle places seemingly diverse phenomena, such as gain modulation, changes in temporal filtering, or network interactions during locomotion, within a common framework (Fig. 1a).

To characterize how natural sensory input changes with locomotion, we recorded videos by moving a camera in different natural environments (Fig. 1b). The camera was held close to the ground, mimicking the visual experience of a rodent, while we performed recordings in two locomotion states: stationary, where the camera moved slightly while being held in place, and during movement (see Methods for details). To understand how locomotion impacts the statistics of natural stimuli and, consequently, signals encoded by visual neurons, we processed videos with linear Gabor filters reminiscent of receptive fields in the mouse primary visual cortex (*16*) (Fig. 1c). We found that the transition from a stationary state to movement substantially changes the filter output, even if both recordings were performed in the same natural scene. The impact of locomotion extends over multiple statistical features: increased variance and dynamic range of filter outputs (Fig. 1d, left), changes in their frequency spectrum (Fig. 1d, middle), as well as decreased magnitude of correlations between filter outputs (Fig. 1d, right).

While our camera recordings capture the full complexity of natural scenes, they coarsely approximate animal movement. In addition to locomotion, freely moving mice move their head and eyes, which adds complexity to the sensory input. To confirm that recordings of natural scenes approximate the real animal experience sufficiently well, we reanalyzed videos recorded from the perspective of freely moving mice, made available by Parker and colleagues (*15*) (Fig. 1e). These videos were recorded using a head-mounted camera as mice freely explored an experimental setup where artificial stimuli were displayed. The resulting recordings were therefore an approximation of the input to the mouse visual system during unconstrained, natural behavior. The dependence of these stimuli on the locomotion state was closely reminiscent of that of the natural scene recordings (Fig. 1f). We highlight this correspondence in different contexts throughout the remainder of this work. More fine-grained behaviors such as eye movements also influence stimulus statistics. However, their effect is likely more subtle than changes caused by locomotion. To confirm this intuition we performed additional analyses on natural scene recordings with simulated saccadic movements as well as videos corrected for the eye position of the mice. We found no qualitative differences in the impact of locomotion on the analyzed stimulus statistics (see Fig. S1 and Fig. S2).

Another important factor to consider is that mice perceive both green and ultraviolet (UV) light. Since the statistics of spectral content vary systematically with elevation (*17, 18*), we investigated how color and elevation influence the effects of locomotion on stimulus statistics. We found that locomotion reliably increases the dynamic range of filter outputs across colors and elevations (see Fig. S3).

To better understand why movement impacts stimulus statistics and how universal this impact is, we constructed a minimal model of sensing during locomotion (Fig. 1g). In the model, a point agent equipped with two sensors moves in a two-dimensional environment in which the average stimulus value changes with spatial position. During a stationary state, the agent’s position fluctuates around a small area, thus sampling a small region of the spatially correlated environment (Fig. 1g, blue frame). In turn, during locomotion, the agent samples larger regions during the same time period. The transition between these behavioral states results in a change in stimuli registered by the agent’s sensors that qualitatively mimics changes in V1-like filter outputs to natural videos and experimental stimuli in freely moving mice (Fig. 1h). Simulations of the minimal model suggest that the observed dependence of stimulus statistics on movement is universal and originates from the coarse spatial features of the scene. It should therefore be experienced by any organism moving through a heterogeneous environment.

### Gain modulation maintains accuracy of sensory coding during locomotion

Perhaps the most salient type of modulation of sensory neurons by locomotion is the increase in gain of sensory responses (*4*). We postulate that the gain increases to match an increase in the stimulus variance experienced during movement (Fig. 2a). To illustrate this effect, we consider a linear Gabor filter, reminiscent of a mouse V1 receptive field (Fig. 2b, inset). The variance of the filter response to videos recorded during camera movement is indeed visibly higher than in the stationary state (Fig. 2b, left panel, orange and gray lines, respectively). This increase is reflected in the distributions of standard deviations of the filter outputs (Fig. 2b, middle panel). Sensory input statistics in freely moving mice, as well as the simulations of the minimal model, are qualitatively in agreement with the statistics of natural stimuli (Fig. 2b, middle panel, inset), suggesting that the increase in variance is a fundamental property of sensory streams experienced during movement in spatially inhomogeneous environments. The variance change observed during locomotion is regular, systematic and it generalizes across different filters, natural videos, and timescales of the analysis (Fig. 2b, right panel). It is also generated by spatiotemporal filters that mimic temporal selectivity of cortical neurons (Fig. S4).

**Figure 2:**
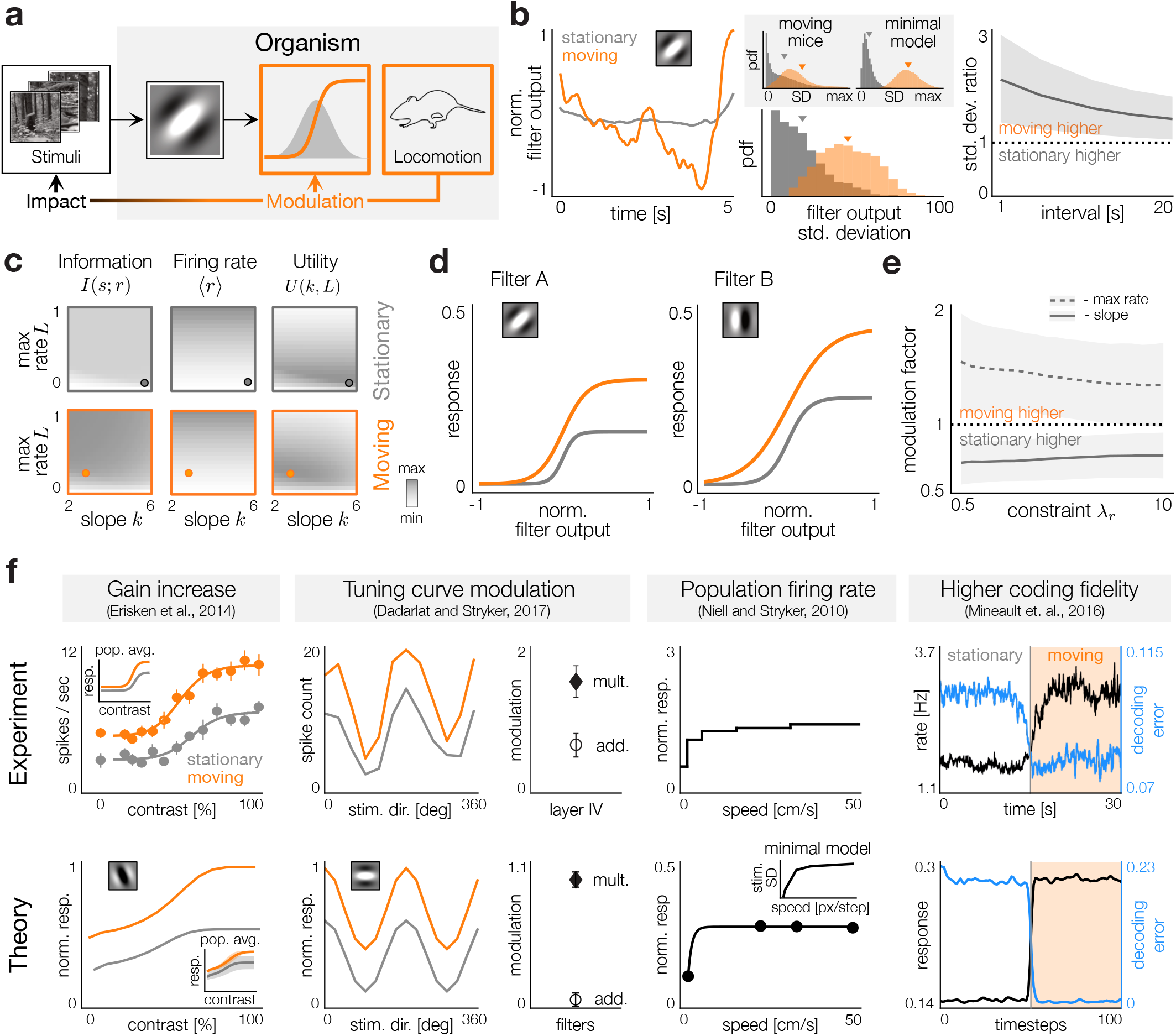
Locomotion impacts stimulus variance and modulates the gain of sensory neurons. **a)** Locomotion impacts filter output variances and modulates the dynamic range of neural nonlinearities. **b)** Left: example filter outputs during locomotion (orange) and stationary periods (gray). Middle: distributions of standard deviations of the example filter across videos in movement (orange) and stationary (gray) conditions. Triangles denote averages. Inset: same distributions obtained from recordings in moving mice (*15*) and the minimal model. Right: ratio of standard deviations of filter outputs during movement to those during stationary periods over different time intervals. Line depicts average and shaded area standard deviation. **c)** Left: mutual information between inputs and neural response *I*(*s*; *r*) as a function of slope *k* and maximum firing rate *L* during movement (top) and stationary state (bottom). Middle: average firing rate. Right: utility function computed for the constraint *λ*_*r*_ = 5. **d)** Example nonlinearities optimized for stationary (gray) and moving (orange) conditions. Insets depict receptive fields of optimized neurons. **e)** Ratio of optimal maximal firing rate *L* (dashed line) and slope *k* (solid line) in the moving condition versus the stationary condition for different constraints *λ*_*r*_ . Lines depict averages and shaded areas standard deviations across the neural population. **f)** Comparisons of published experimental results (top) and theoretical predictions (bottom). From the left: responses of an example neuron and the population (inset) to visual contrast (*5*), modulation of orientation tuning curves (black and white symbols denote population average multiplicative and additive modulation respectively, bars denote standard deviations) (*8*), population firing rate as a function of speed (*4*) (bottom: circles - average responses to videos recorded at different speeds, black line - exponential fit, inset - results from the minimal model), decrease of decoding error (blue) and increase in firing rate (black) at the transition to movement (*13*).

To efficiently and accurately encode stimuli whose variance increases rapidly, sensory neurons must expand their capacity to match the change in stimulus statistics (*19–22*). The systematic increase in stimulus variance with locomotion suggests a modulation strategy in which sensory neurons increase their dynamic range to accommodate the greater variability of the stimulus received during movement. However, expanding the dynamic range while keeping the maximal firing rate constant inevitably lowers the fidelity of the sensory code, as broader stimulus distributions are mapped onto the same limited range of responses. To rigorously test this intuition, we constructed simple linear-nonlinear models of individual neurons, where the output of each Gabor filter is transformed to simulated neural activity through a logistic nonlinearity (Fig. 2a). For each of the locomotion states, we optimized the slope term *k* and the maximal firing rate *L* to maximize the utility function *U*(*k, L*) that trades off information transmission, measured as the mutual information *I(s; r*) between filter outputs *s* and neural activity *r*, with the average firing rate ⟨*r*⟩, subject to a constraint λ_*r*_ (Fig. 2c, see Methods for details). To focus only on changes in the dynamic range of stimuli, we assumed that neurons adapt to the mean change at the timescale of the considered excerpts. We found that, in order to match the systematic increase in stimulus variance, neural nonlinearities optimized for the locomotion state simultaneously decrease their slope *k* to expand their dynamic range, and increase their maximal firing rate *L*, compared to the stationary state, to maintain high-fidelity information transmission (Fig. 2d, orange and gray lines, respectively). This indicates that the benefit of encoding higher-variance stimuli with high fidelity during locomotion outweighs the metabolic cost of a higher average firing rate. These changes generalize over a range of constraint values λ_*r*_ and across a broad range of model Gabor receptive fields (Fig. 2e). The predicted increase in gain during movement is also observed in neuron models that include input and output noise (Fig. S5).

The relatively simple modulation of neural nonlinearities optimized to efficiently encode stimuli whose variance is impacted by locomotion can already account for a range of experimental results published previously (Fig. 2f). The expansion of nonlinearities explains the increase in the gain of contrast tuning curves during locomotion reported in (*5*) (Fig. 2f, left panel) and the modulation of orientation tuning curves reported in (*8*) (Fig. 2f, second from the left). Beyond individual neurons, the theory provides a quantitative match at a population level, where locomotion-induced modulation of orientation tuning curves exerts a stronger multiplicative rather than additive effect (Fig. 2f, second from the left, black and white symbols, respectively). To understand how locomotion at different speeds should modulate neuronal gain, we optimized model neurons to statistics of natural videos recorded at different speeds of movement. Interestingly, optimal gain modulation does not continue to increase with locomotion speed but saturates quickly after the onset of movement, which is reflected in the simulated population activity (Fig. 2f, third from the left, bottom), as well as the stimulus statistics in the minimal model (Fig. 2f, third from the left, bottom, inset). This result matches the experimental findings of (*4*) (Fig. 2f, third from the left, top), suggesting that there is no benefit in expanding the dynamic range of neurons beyond what is required to accurately encode stimuli observed at relatively low movement velocities. This result is robust to the statistics of individual environments and different spatial frequencies of the Gabor filters (Fig. S6). Finally, the increased dynamic range of sensory neurons during locomotion results in improved accuracy of sensory coding if the dynamic range of the stimulus is kept constant, as in the experiment. This improvement has been reported by (*13*), who quantified it using the population decoding error (Fig. 2f, right panel, top). We performed an analogous analysis on simulated neurons and found that the population decoding error decreases at the transition to locomotion, which is accompanied by an increase in the average firing rate, thus resembling experimental results (*13*) (Fig. 2f, right panel, bottom row, blue and black lines, respectively).

### Modulation of temporal filtering ensures efficiency of sensory coding during locomotion

Beyond gain increases, locomotion alters temporal processing and induces more sustained responses (*6*). We propose that this form of modulation adjusts sensory neurons to changes in the temporal statistics of stimuli induced by locomotion (Fig. 3a). These changes are apparent in the average temporal spectra of the response of the Gabor filters to natural videos, where locomotion causes an increase in the spectral power at all temporal frequencies relative to the stationary condition (Fig. 3b, left panel). During movement, a sensory neuron with a Gabor-like spatial receptive field is subject not only to a substantial change in the total power of its input (as reflected by variance; see previous section), but also to a change in its frequency content. Statistics of videos from the perspective of freely moving mice, as well as the minimal model, reproduce the observed increase in spectral power seen in natural stimuli (Fig. 3b, left panel, inset). This correspondence supports the hypothesis that such temporal dynamics are intrinsic to sensory experiences during locomotion through spatially diverse settings.

**Figure 3:**
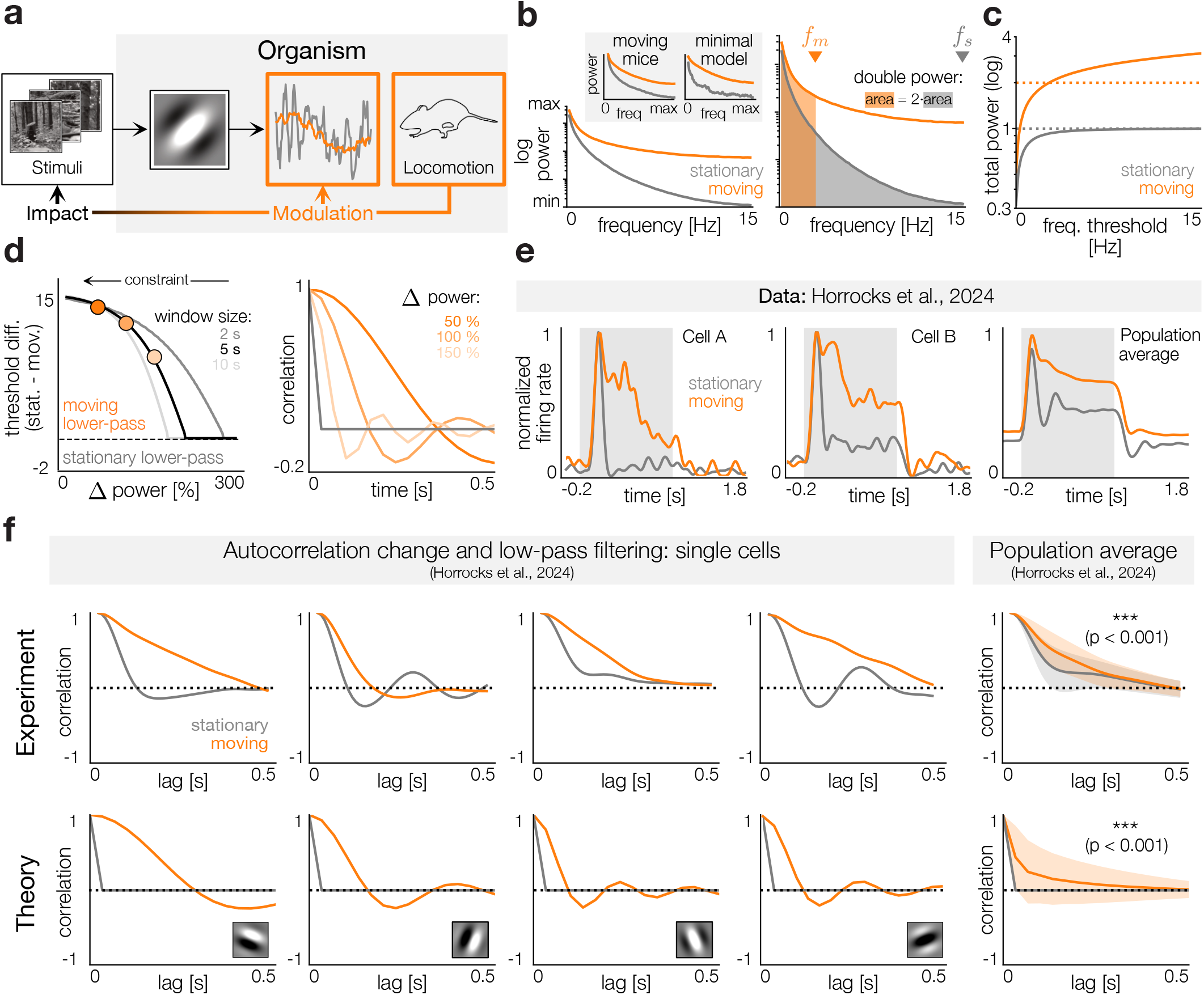
Locomotion impacts stimulus frequency spectrum and modulates temporal filtering in sensory neurons. **a)** Locomotion impacts the temporal spectra of filter outputs and modulates temporal filtering in individual neurons. **b)** Left: the average power spectrum of filter outputs during locomotion (orange) and rest (gray). Inset: same spectra from recordings in moving mice (*15*) and the minimal model. Right: a temporal filter with a frequency threshold transmits a fraction of the signal power computed as the area under the curve. Frequency threshold during movement *f*_*m*_ and stationary condition *f*_*s*_ are denoted by orange and gray triangles, respectively. The total power during movement (orange area) is twice that of the stationary condition (gray area). **c)** The total power transmitted by the temporal filter as a function of the frequency threshold for stationary (gray) and moving conditions (orange). Dashed lines: example power levels considered in (b). **d)** Left: The difference in the frequency threshold between the stationary and moving conditions for a range of energy constraints (Δ power). Right: The autocorrelation functions of temporal filters for the stationary (gray) and moving conditions (orange). Shades of orange correspond to energy constraints denoted with orange circles in the left panel. **e)** The responses of mouse V1 neurons to moving dot stimuli (gray background indicates stimulus period) during locomotion (orange) and stationary (gray) states. Experimental data from (*6*) is reanalyzed under [CC BY 4.0](https://creativecommons.org/licenses/by/4.0/). **f)** Comparisons of reanalyzed experimental results from (*6*) (top) and theoretical predictions (bottom). Top: The autocorrelation functions of trial-averaged responses of single neurons. Top right: population averages for the stationary and moving conditions. Shaded areas denote standard deviations. The decay time is significantly longer in the moving condition. Bottom: Autocorrelations of individual model neurons whose temporal filters correspond to the 50% increase energy constraint. Bottom right: Same as top-right but for population of model neurons. Top panels plotted from data provided by (*6*) under [CC BY 4.0](https://creativecommons.org/licenses/by/4.0/).

Sensory neurons operate under metabolic constraints and must therefore balance their activity with the need to transmit information. In the temporal domain, this entails prioritizing frequencies with higher power, because they contain a larger fraction of stimulus information (*11, 23*). Ideally, a neuron would encode the same frequency range in all behavioral states. However, the elevated power spectrum during locomotion poses a challenge: even with increased metabolic capacity, a neuron may not be able to transmit all frequencies and instead needs to prioritize those with higher power.

To formalize this idea, we modeled the temporal processing of neurons as a linear filter in the frequency domain that transmits frequencies below a certain threshold (including a linear roll-off; see Methods), reflecting the power-law decay of temporal frequency power observed in natural stimuli (Fig. 3b). We considered a scenario where, rather than being fixed, the frequency threshold adapts to the changes in temporal statistics of stimuli by increasing the metabolic resources available during locomotion. This adaptive threshold modulated by the locomotion state can therefore take two different values *f*_*m*_ and *f*_*s*_ during movement and stationary states (Fig. 3b, right panel, orange and gray triangles, respectively). For example, when the frequency threshold for the stationary state enables transmission of all frequencies (i.e. *f*_*s*_ = 15 Hz), the frequency threshold *f*_*m*_ that corresponds to a 100% increase in total power with respect to the stationary condition is equal to 2.9 Hz (Fig. 3b, right panel, shaded areas), demonstrating the prioritization of lower temporal frequencies which have higher power. To systematically assess the effect of changing the frequency threshold, we computed the total power transmitted through the temporal filter as a function of frequency threshold in stationary and moving conditions (Fig. 3c). Using the power required to transmit the entire spectrum in the stationary condition as a reference for neuronal capacity, we systematically increased the allowable power during movement and determined the corresponding frequency thresholds. Across a wide range of metabolic constraints computed as a percentage increase in power relative to the stationary state, the filter thresholds in the moving condition were consistently lower than during rest (Fig. 3d, left panel). This effect is further evident in the autocorrelation functions of the temporal filter responses to broadband white noise signals (Fig. 3d, right panel). In the stationary state, all frequencies are passed through by a model neuronal filter, and the autocorrelation of the output is equal to that of the white noise input, i.e. a delta function (Fig. 3d, right panel, gray line). During locomotion, the model neuron acts as a low-pass filter, which is reflected in a slower decay of the corresponding autocorrelation functions, where the decay time is determined by the strength of the metabolic constraint (Fig. 3d, right panel, orange lines). These analyses highlight that even when metabolic constraints are relaxed, temporal processing during locomotion should remain more biased toward lower frequencies than during stationary periods. The theory thus generates a clear prediction: neurons should exhibit more pronounced low-pass temporal filtering during locomotion than during rest.

To test this prediction, we reanalyzed data from Horrocks et al. (*6*), who recorded neuronal responses in mouse V1 to moving dot stimuli during periods of locomotion and rest (Fig. 3e). We computed autocorrelation functions of the average neural responses in stationary and moving conditions, as well as the average autocorrelation across the population (Fig. 3f, top row). Comparison of empirical autocorrelations with theoretical predictions for individual Gabor filters and the filterbank average revealed close agreement between data and theory (Fig. 3f, bottom row). As a quantitative metric, we calculated the decay time (the time lag at which the autocorrelation drops to 0.5) and found it was significantly longer during locomotion in both the experimental data and a population of simulated model neurons (Fig. 3f). We note that in the experimental setting considered here the sensory input is held constant across behavioral states. However, in natural conditions, the input statistics change significantly during movement, and the power of higher frequencies increases strongly. Therefore, it is possible that despite low-pass filtering, neural activity in response to natural sensory input is less temporally correlated during movement than at rest. Together, these results demonstrate that the temporal processing strategies predicted by the proposed efficient coding principle are reflected in neural responses, supporting the idea that locomotion modulates temporal filtering in V1 to account for systematic changes in temporal stimulus statistics.

### Modulation of network interactions maintains a non-redundant sensory code during locomotion

Locomotion modulates not only individual neurons, but also activity at the population level by modifying interactions between cells (*8*). A prominent manifestation of this effect is changes in the spatial integration of visual stimuli during locomotion, which is likely mediated by network interactions modulated by the behavioral state (*14*).

As in the previous sections, our first aim was to characterize how inputs to multiple neurons change simultaneously during locomotion. To do so, we analyzed the correlations of responses of receptive-field-like filters to natural videos (Fig. 4a). To ensure that filters were non-redundant and optimized as a population, we learned them using independent component analysis (ICA), which reproduces the properties of V1 receptive fields (*24, 25*) (Fig. 4c, right panel). ICA was trained on a large set of image patches consistent with the size of the receptive fields in mouse V1 (Fig. 4c, left panel). We observed that locomotion has a strong impact on the correlations of the filter outputs. In stationary periods, filter responses tend to be highly redundant due to the shared, slowly changing sensory drive corresponding to slow fluctuations of the visual field (Fig. 4b, left panel, gray lines). The onset of locomotion significantly decreases the magnitude of filter correlations (Fig. 4b, middle panel, orange lines). As previously, we confirmed that this pattern is systematic and that the magnitude of the correlation is significantly stronger during the stationary state across all filter pairs and natural videos, as well as videos recorded in behaving animals (Fig. 4b, right panel). The minimal model matches the statistics of the natural videos; neighboring sensors of an agent moving in two-dimensional environments reveal a similar dependence of correlation magnitude on locomotion speed, suggesting the universality of this pattern.

**Figure 4:**
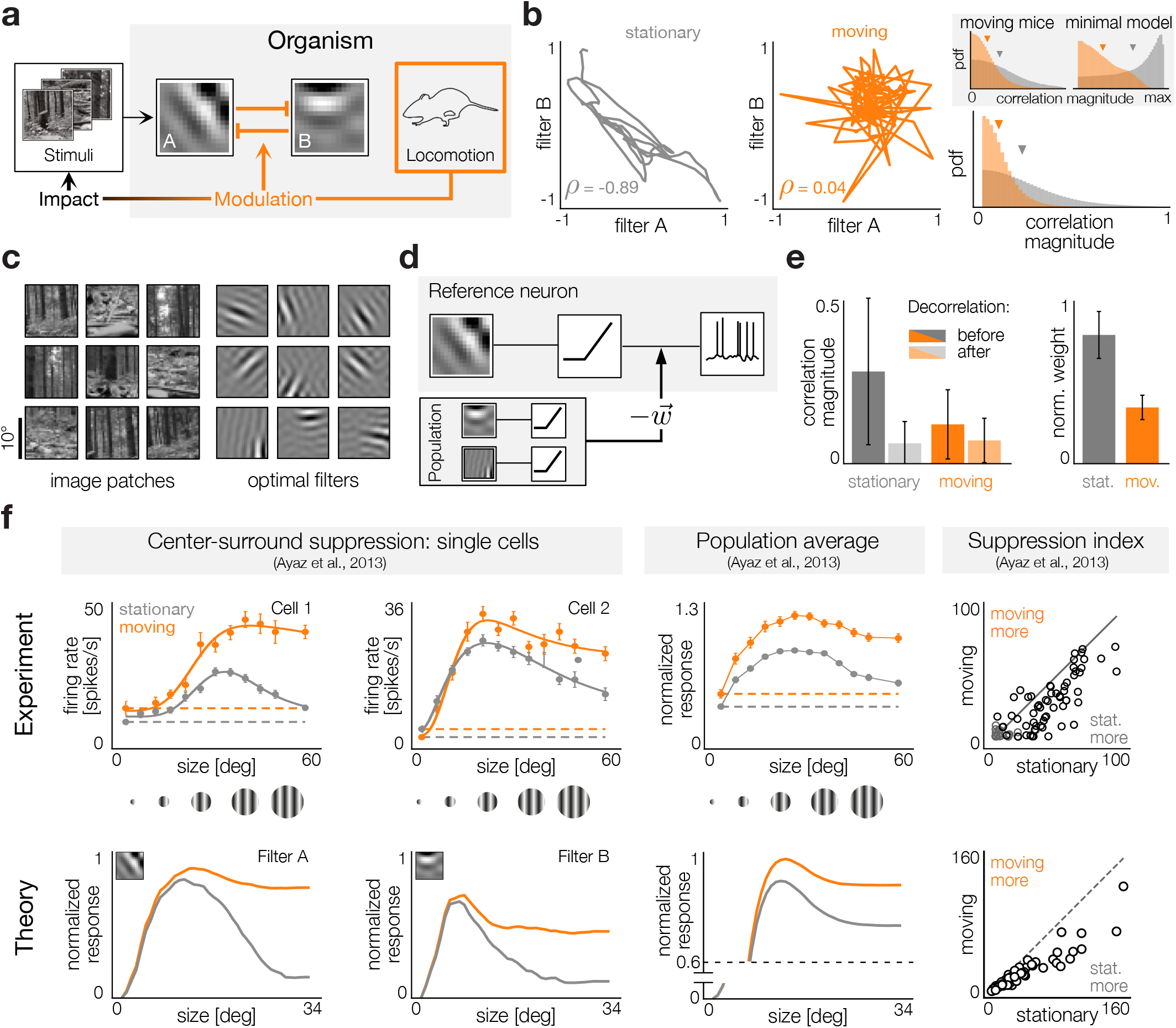
Locomotion impacts filter correlations and modulates lateral inhibition in sensory populations. **a)** Locomotion impacts the correlation structure between filter outputs and modulates network interactions. **b)** Left and middle: the correlation of the outputs of two filters in five second windows for the stationary (gray) and moving (orange) conditions. Right: distribution of correlation magnitudes across many windows in the stationary (gray) and moving (orange) conditions. Triangles indicate averages. Inset: same magnitudes from recordings in moving mice (*15*) and the minimal model. **c)** Left: example natural image patches. Right: example independent component filters. **d)** Schematic of lateral inhibition in a network of model neurons. The outputs of the population are rectified, weighted, and subtracted from the rectified output of a reference model neuron. **e)** Left: average magnitude of correlation between model neuron pairs in stationary (gray) and moving (orange) conditions before (darker) and after (lighter) decorrelation. Right: optimal weights (normalized such that the vertical axis limit is unity) for the stationary (gray) and moving (orange) conditions. Error bars denote standard deviations. **f)** Comparisons of published experimental results (top) and theoretical predictions (bottom). Experimental results were replotted from (*14*) under CC BY 3.0. Responses for the stationary and moving conditions are in gray and orange respectively. First from the left: individual V1 neural (top) and model (bottom) responses to oriented grating stimuli of increasing size. Middle: population response for 60 neurons (top) and 101 model neurons (bottom). Right: the suppression index of individual neurons (top) or model neurons (bottom).

Strong correlations between filters are a form of redundancy. Redundant neurons simultaneously convey the same information, which decreases the efficiency of sensory coding in the low-noise regime (*26*). To minimize redundancy, neuronal networks should decorrelate their inputs (*26, 27* ). In the visual cortex, center-surround interactions are believed to reflect lateral inhibitory interactions within a local network that support efficient neural representations through redundancy reduction (*28, 29*). These observations suggest a strategy for modulation of neuronal networks by locomotion state. During stationary periods, correlations between receptive fields are high and neurons should decorrelate their outputs, which is achieved by strong lateral inhibition. During locomotion, correlations between sensory inputs decrease, hence inhibitory interactions should weaken since there is no need to decorrelate stimuli. This network modulation mechanism ensures coding efficiency across behavioral states at the level of neuronal populations.

Following these lines of thought, we developed a model of lateral inhibition (Fig. 4d). Our approach is based on previous models of redundancy removal in natural scenes through local interactions in a sensory population (*29*). In the model, the output of each linear receptive field was transformed with a thresholding nonlinearity to a positive firing rate. The activity of each neuron was linearly suppressed by other neurons in the population to minimize the magnitude of correlations (i.e. redundancy; see Methods for details). For each neuron and each stimulus interval, this lateral inhibition was controlled by an independently optimized synaptic weight vector 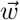. The lateral inhibition model successfully reduces the correlations between all pairs of model neurons and during all stimulus intervals (Fig. 4e, left panel). The decrease is most prominent during the stationary state, when the initial correlation magnitude is high (Fig. 4e, left panel, gray bars). However, even during locomotion when the correlation magnitude is much weaker, the relative effect of decorrelation is also noticeable (Fig. 4e, left panel, orange bars). We note that in reality, sensory systems must dynamically adapt their decorrelation mechanisms to changing stimulus statistics, a process which we approximate by optimizing weights independently for individual stimulus excerpts of 5 seconds. Although the neural implementation of dynamic decorrelation goes beyond the scope of this work, we believe that it could be instantiated using mechanisms similar to decorrelation (*30*) or adaptive whitening by inhibitory neurons (*31*).

The lateral inhibition model fit to natural videos predicts that, to maintain high efficiency of coding, the state of the network should adjust by decreasing the magnitudes of inhibitory interactions during locomotion (Fig. 4e, right panel). To verify these predictions, we simulated neural responses by exposing model neurons to oriented sinusoidal gratings of increasing diameter, a set of stimuli used to test spatial integration during locomotion (*14*) (Fig. 4f). To simulate different states of movement, we used two sets of inhibitory weights optimized for one excerpt of the natural video during the stationary state and one during locomotion. When the diameter of the stimulus extends beyond the receptive field of a neuron, it activates other members of the population, thus inducing an inhibitory effect and suppressing the response of the analyzed neuron. Due to differences in the optimized weights, suppression is much stronger in the simulated stationary condition than during movement (Fig. 4f, bottom row, first and second panels from the left, gray and orange lines, respectively). This effect matches the results reported by (*14*) at the level of individual cells (Fig. 4f, top row, first and second column from the left), as well as the population average computed across all model neurons, and natural video excerpts (Fig. 4f, third column from the left). The inhibitory effect can be further quantified by the suppression index (*14*), whose values are lower during locomotion and qualitatively match the experimental results (Fig. 4f, right column). A close resemblance of simulations and experimental data suggests that network interactions can also be modulated by locomotion to optimize processing of correlated stimuli in the visual cortex.

### Stimulus statistics explain differential modulation of sensory coding by locomotion in rodents and primates

An intriguing recent line of research revealed that modulation of sensory areas by locomotion is not universally preserved in the animal kingdom. In contrast to rodents, movement has a very weak effect on sensory responses in V1 of marmoset monkeys (*9*). It is unclear whether this disparity implies different species-specific computational objectives or whether it could be understood within the same theoretical framework.

An important difference between receptive fields of neurons in mouse and marmoset V1 is their size and spatial frequency. While mouse neurons have large, low-frequency receptive fields on the order of 10-20 degrees of visual angle (*16*), the foveal receptive fields in monkeys are much smaller and higher-frequency (*9, 32, 33*). To understand the impact of locomotion on higher spatial frequencies, we analyzed responses of Gabor filters of differing sizes and spatial frequencies (Fig. 5a). As we have shown in previous sections (see Fig. 1 and Fig. 2), the outputs of a representative mouse-like receptive field (Fig. 5a, left panel, blue frames) fluctuate much more during movement than during stationary periods (Fig. 5a, middle panel, orange and gray lines, respectively). The output of a significantly smaller, higher frequency Gabor filter reminiscent of a primate foveal receptive field (Fig. 5a, left panel, yellow frames) behaves differently. Already in the stationary state where the field of view moves slowly, the filter output fluctuates very rapidly, and the relative change in the filter output variance during movement is negligible (Fig. 5a, right panel). To establish whether this effect is systematic and may be the basis for the previously observed differences between species, we generated two sets of Gabor filters whose spatial frequencies were matched to receptive fields recently measured in mouse and marmoset V1 reported in (*32*) (Fig. 5b). The standard deviations of mouse-like filter responses to videos recorded during movement are on average almost twice as high as those for stationary videos (Fig. 5c, left panel, blue bar). The relative change in standard deviations of marmoset-like filters between stationary and moving conditions is minimal, and their average ratio is 1.14 (Fig. 5c, left panel, yellow bar). We emphasize that while the standard deviations of marmoset-like filter outputs have lower magnitude than the standard deviations of mouse-like filter outputs due to the statistics of natural scenes, the predictions about gain modulation depend only on the *relative* change in filter output standard deviations between stationary and moving conditions. We also note that the variance of filter modulation is relatively high for low-spatial-frequency filters and it practically vanishes for high spatial frequencies (Fig. 5c, left panel, error bars).

**Figure 5:**
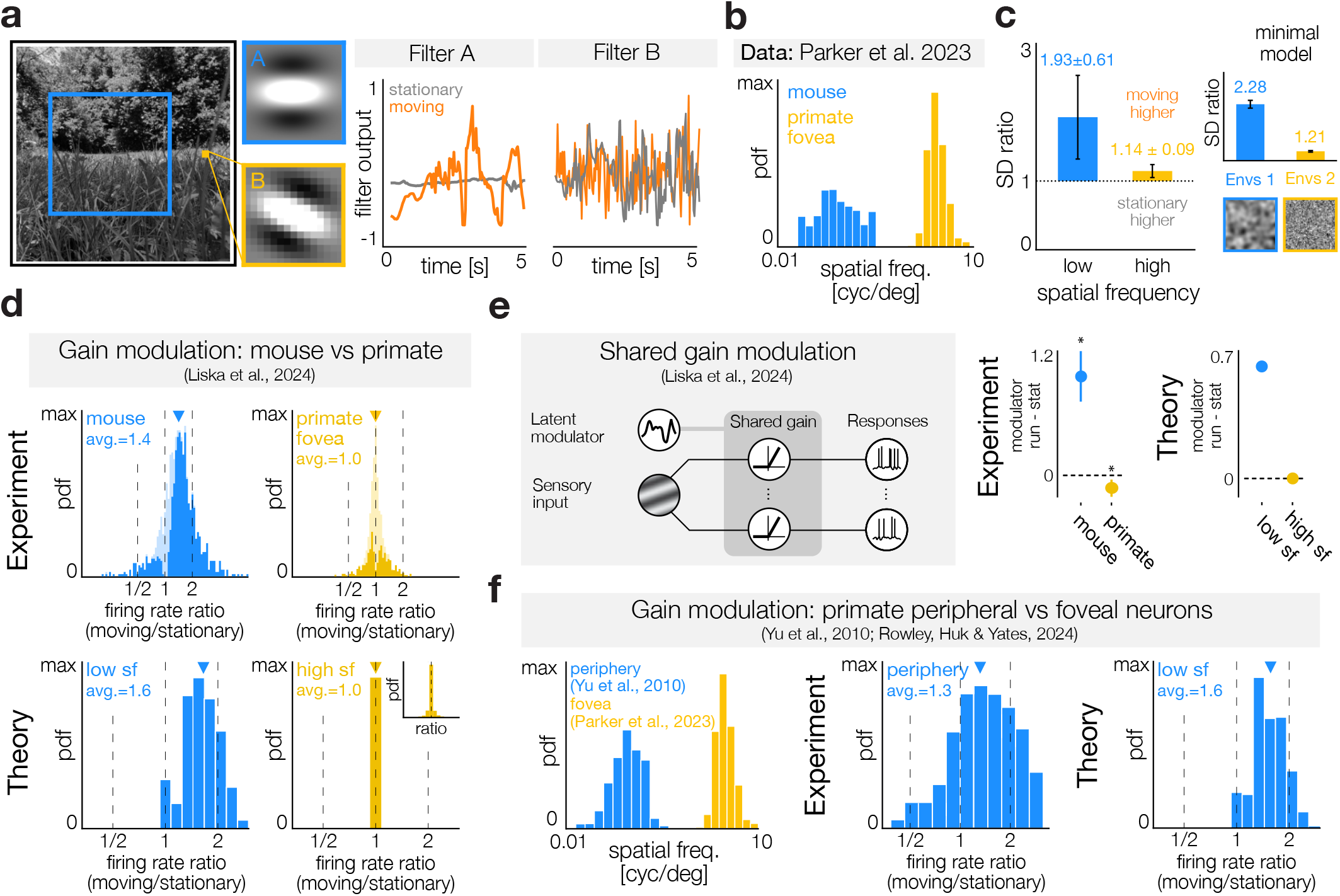
Locomotion differentially impacts high and low spatial frequency filters and modulates the gain of sensory neurons in different species. **a)** Left: Gabor filters with a low (blue; 0.05 cyc/deg) and high (yellow; 2 cyc/deg) spatial frequency. Middle and right: outputs of the low- (middle) and high- (right) frequency filters in a stationary (gray) and moving (orange) conditions. **b)** Distributions of spatial frequencies of receptive fields in mouse V1 (blue) and foveal region of the marmoset V1 (yellow). Data from (*32*). **c)** Ratios of standard deviations (SDs) of filter responses during movement to SDs during stationary periods. Left: SD ratios of filters whose spatial frequencies were matched to mouse V1 (blue) and foveal V1 in the marmoset (yellow). Error bars indicate the SD. Right: analogous analysis in the minimal model in environments with large (blue) and small (yellow) spatial scales (illustrated at the bottom). **d)** Top: distributions of the average ratio of firing rates of V1 neurons recorded during stationary periods and movement in the mouse (left) and marmoset (right). Replotted from (*9*) under [CC BY 4.0](https://creativecommons.org/licenses/by/4.0/). Bottom: analogous to the top row but for simulated neurons with receptive fields matched in spatial frequency to the mouse (blue) and marmoset (yellow) V1. **e)** Left: scheme of a shared gain modulator model. Middle: difference between the average values of the shared modulator during locomotion and stationary periods in the mouse (blue) and the marmoset V1 (yellow), replotted from (*9*) under [CC BY 4.0](https://creativecommons.org/licenses/by/4.0/). Right: analogous to the middle panel but for simulated neurons (see Methods for details). **f)** Left: distributions of spatial frequencies of peripheral (blue) and foveal (yellow) receptive fields in the marmoset V1. Data from (*33*) and (*32*) respectively. Middle: analogous to panels in (d), but for neurons with peripheral receptive fields in the marmoset V1. Replotted from (*34*) under CC BY-NC-ND 4.0. Right: analogous to the middle panel but for simulated neurons with receptive fields matched in spatial frequency to the peripheral receptive fields in marmoset V1.

These disparities between mouse- and primate-like filter outputs can be explained by a minimal model of sensory signals during movement (Fig. 5c, right panel). In a spatially correlated environment, locomotion increases the variability of the sensor output (Fig. 5c, right panel, blue frame and bar, respectively). In natural scenes, such spatially correlated inputs are processed by low-spatial-frequency filters. In contrast, high-frequency filters process visual features that are much less spatially correlated. Movement in a simulated environment that matches this property and is weakly correlated does not impact the sensor variance (Fig. 5c, right panel, yellow frame and bar, respectively). Minimal model simulations suggest therefore a universal relationship between the spatial frequency of the filter and the impact of locomotion.

Our next step was to understand how the frequency-specific impact of locomotion on filter output should affect the gain of sensory neurons. Following the procedure described in a previous section (see Fig. 2), we optimized nonlinearities in two populations of model neurons with mouse- and marmoset-like receptive fields. To emulate experiments comparing these two species (*9*), we exposed these two populations of model neurons to identical random stimuli (see Methods). We simulated transitions from the stationary state to running by switching nonlinearities optimized to natural videos recorded in these two conditions. Following the analysis of (*9*), we quantified the locomotion-induced gain change by computing the ratio of the firing rates during locomotion and the stationary state. The distribution of the firing rate ratios of model neurons closely resembles V1 neurons in mice and marmosets (Fig. 5d). In mice and low-frequency model neurons, the average activity is significantly increased during locomotion (Fig. 5d, left column, blue). We observe a quantitative match between the average ratio of firing rates during running and rest (the average ratio reported in (*9*) is 1.402, 95% confidence interval (CI) [1.365, 1.440], while we find that the average ratio for the low frequency model neurons is 1.65, 95% CI [1.63, 1.67]). In contrast, the marmoset and high-frequency model neurons are not modulated on average (Fig. 5d, right column, yellow). In both experimental data and theoretical predictions, the average firing rate ratio is equal to 1 (the average ratio reported in (*9*) is 1.011, 95% CI [0.995, 1.027], while we find that the average ratio for the high frequency model neurons is 1.00002, 95% CI [0.99996, 1.00008]). The match between theory and data goes beyond averages of firing rate ratios—the variability of modulation by locomotion is higher in mice than in marmosets, as quantified by the confidence intervals. The experimental study (*9*) has reported that the gain of foveal marmoset neurons can be minimally suppressed by locomotion, when encoding stimuli with preferred orientation, but remains unchanged when considering all stimuli. We believe that the weak suppression to preferred stimulus orientations may be caused by factors that go beyond the statistics of natural stimuli analyzed here. Our theory provides an explanation of the behavior of neurons averaged across a broad range of stimuli encountered in natural environments.

The transition to movement modulates the gain of many neurons simultaneously. This population-level effect can be quantified using a shared modulator model that infers a single latent variable that controls the gains of neural responses independently of the sensory input (*9*) (Fig. 5e, left panel). In mice, the onset of movement causes a strong increase in the response gain, which is reflected in the difference between the average modulator values during running and rest (Fig. 5e, middle panel, plot reproduced from (*9*)). In marmosets, this difference becomes weakly negative and effectively vanishes (Fig. 5e, middle panel, plot reproduced from (*9*)). We performed an analogous analysis on simulated populations by approximating the shared gain modulator using linear methods (see Methods). In mouse-like model neurons, the increase in firing rates induced by the onset of locomotion can be captured by a single latent modulator, which dramatically changes its value between behavioral states (Fig. 5e, right panel). In marmoset-like neurons, the transition between locomotion states does not cause a population-level gain change; therefore, the difference between modulator values is effectively zero (Fig. 5e, right panel). We therefore conclude that modulation which preserves the efficiency of the sensory code offers an explanation of population-level effects associated with locomotion in these two species.

The results described above suggest that the differences between the modulation of V1 in mice and marmosets are caused by changes in input statistics associated with locomotion and do not result from species-specific computational principles. This observation generates an immediate prediction: even in the same species, locomotion-induced gain modulation should be larger in neurons tuned to low spatial frequencies, whereas neurons with high-frequency receptive fields should remain unaffected. A recent study has shown that peripheral neurons in marmoset V1, whose receptive fields are comparable to mouse neurons, are significantly modulated by running (the average ratio of the firing rates reported in (*34*) is 1.33). Following the procedure outlined above, we simulated a population of neurons with peripheral-like receptive fields (Fig. 5f, left and right panels; we relied on the data published in (*33*)). The gain changes quantified by firing rate ratios in the simulated neurons are in agreement with the experimentally measured activity of marmoset neurons with peripheral receptive fields (Fig. 5f, right and middle panels, respectively; the average ratio is 1.57, 95% CI [1.55, 1.59]). Our findings thus suggest that differences between mice and marmosets are a manifestation of the same computational principle—modulation that adjusts sensory neurons to systematic changes in stimulus statistics evoked by behavior.

## DISCUSSION

The simple yet general principle of adapting to the statistical structure of sensory input can explain many seemingly disparate effects that locomotion exerts on sensory systems. By extending the efficient coding hypothesis, we provide a unifying account of phenomena observed within individual species: modulation of gain, changes in temporal response dynamics, and spatial integration in mice, as well as differential modulation of foveal and peripheral neurons in primates. The theory was also able to reconcile the differences between species in gain modulation and the dimensionality of population responses during locomotion. We thus demonstrate that differences in locomotion-induced modulation of sensory systems do not need to be a species-specific computation, but a manifestation of a general coding principle.

### Towards a theory of sensory coding in behaving animals

Theories that underlie our understanding of sensory systems are largely concerned with passive processing and transmission of sensory information (*12, 35*). In its original statement, the efficient coding framework focuses on properties of natural stimuli (*10, 25*) and does not reference the interplay between the behavior of the organism and statistics of the sensory experience. In this work, we make a step towards developing a theory of sensory coding in freely behaving organisms. Among the many ways in which behavior may impact sensory input we focus on a relatively coarse, yet prominent and relevant, behavioral state—locomotion. Our key proposal is that the strong and predictable change in stimulus statistics caused by the onset of movement is a form of regularity that can be exploited by the nervous system in the same way that receptive fields of neurons adapt to the correlation structure of natural scenes to increase efficiency of sensory codes (*25, 36, 37* ). However, in contrast to tuning properties of sensory neurons, this form of adaptation to the sensory niche needs to take the form of an internal modulation mechanism. This modulation dynamically modifies gains, temporal filtering, and network interactions, which improves the coding accuracy and minimizes resource use.

The proposed principle is related to theories of sensory adaptation that are embedded within the efficient coding framework (*12, 19, 20, 38*). Adaptation is a change in a sensory code that occurs in response to a change in the environment in order to maintain the efficiency and accuracy of the representation. The key difference is that changes in the environment occur independently of the organism and have to be estimated from sensory input (*22, 39*–*41*). In contrast, fluctuations in stimulus statistics that are systematically and predictably caused by behavior do not have to be inferred or estimated. The brain can modulate the sensory system using only internal signals to maintain the efficiency of coding during movement and rest.

### Extensions and future directions

A natural direction for future work is to explore progressively better approximations of natural sensory experience in freely behaving animals. One important factor is color vision. Mice perceive two colors: green and ultraviolet (*17* ). The effects described here likely generalize to mouse color vision: the green channel in commercial cameras corresponds to the mouse spectral sensitivity, stimuli in UV and green channels are strongly correlated (*17* ) and UV can be approximated by the blue channel, which is also captured by our recordings (*42*). At a more fundamental level, our minimal model demonstrates that the impact of locomotion on stimulus statistics is a general property of movement in heterogeneous environments. At the same time, color stimuli in natural environments have interesting and nuanced structure. For example, green and UV channels are more strongly correlated in the lower that in the higher visual field (*17, 18*) (although the effects described here persist across the entire visual field - see Fig. S3). Sensory systems exploit these dependencies by instantiating opponent color representations (*43, 44*) and adapting to the differences between upper and lower visual fields (*37, 45*). It is therefore likely that behavior impacts visual stimuli in different colors and parts of the visual field in subtle yet structured ways, and that these changes are exploited by modulatory mechanisms similar to those described here. Future work will reveal how more nuanced, behaviorally induced changes in the ecologically valid range of the light spectrum can be exploited by the mouse visual system (*46*).

Across the animal kingdom, the fine-grained statistics of sensory experience are also affected by species-specific behaviors. In this work, we focus on two very general behavioral states shared by almost all species: locomotion or staying still. Our analyses, as well as the minimal model, show the generality of the impact that the onset of movement has on sensory stimuli. However, the differences in movement of rodents and primates certainly influence the higher-order stimulus features they experience. Understanding the impact of species-specific movements on sensory coding is one of the tantalizing directions for future research (*47* ).

Even within the same species, forward locomotion is only one movement pattern that can systematically change sensory input. We expect that other types of movement associated with systematic changes in stimulus statistics can also modulate neurons to optimize the sensory code. In one example, looking up can increase the response strength of neurons in the mouse visual thalamus (*48*). Because contrast increases with elevation in the natural visual field of a mouse (*17, 18, 37* ), increasing the gain of neural responses during head raises may be an adaptation to maintain the efficiency of the sensory code.

On average, firing rates of neurons in the mouse visual cortex increase at the onset of movement and saturate with increasing running speed (*4*). The theory proposed here predicts this effect based on the changes of stimulus variance at different speeds of movement in heterogeneous environments (Fig. 2f). A fraction of neurons in the mouse visual cortex deviate from this increasing and saturating gain modulation pattern. Running speed can for example decrease gain of sensory responses, or modulate it non-monotonically (*7, 49, 50*). Our theory offers a possible, speculative interpretation of this diversity. Neurons whose gain increases and saturates with running speed may be functioning as “sensors”. In this view, their role would be to encode sensory inputs with high accuracy, and convey them downstream. Neurons with non-saturating gain modulation, could play different roles. Future work could explore and test this hypothesis further.

The computational principles described here may be implemented in many different ways by neural circuits. In one example, decorrelation of stimuli could be achieved either via lateral inhibition or by top-down feedback signaling (*51*). The locomotion state could also impact many more features of neuronal tuning. In particular, coordinated modulation of joint spatiotemporal receptive fields could in principle further improve the efficiency of sensory coding in different behavioral states.

An additional set of questions concerns the interaction between stimulus statistics in different environments and locomotion. Here, we consider systematic, average changes in input statistics elicited by movement in different natural scenes. These relationships could be more nuanced and the activity of sensory neurons is likely modulated by a combination of internal signals, as well as adaptation to scene statistics. Movement in absolute darkness is a particularly salient example of such an interaction: because there is no visual input to encode, behavioral modulation could be in principle suppressed as well. In mice, V1 neurons are modulated during running in total darkness (*4, 7* ). These results are, however, consistent with our interpretation: internal modulation prepares neurons for stimulus statistics experienced *on average* throughout the visual experience, and not in a particular sensory scene (e.g. dark environment).

The influence of behavior on sensory coding takes many different forms (*1*) and extends beyond the effects described here. Although, on average, locomotion increases the strength of sensory responses in mice, it may also decrease it or modulate it in a non-monotonic way (*7, 49*). These modulation patterns can also vary between different cell types (*52, 53*), suggesting a diversity of computational functions and objectives. In addition to locomotion, saccades are another behavioral factor that is particularly important for sensory coding. Eye movement patterns vary between species (*54*) and may play different roles in animals with and without a fovea (e.g. primates and mice, respectively). Recent results show that saccades interact with V1 activity in nuanced ways (*32, 55*), suggesting that the computational objectives of such interactions are different from the principles discussed here. In addition to saccades, cortical activity in mice is driven by fine-scale orofacial movements such as whisking (*56*) as well as “twitching” i.e. minor non-goal-oriented behaviors (*57* ). Interestingly, this kind of modulation is absent in macaques (*47, 58*). Because small body movements are unlikely to dramatically influence the statistics of sensory input, they are unlikely to be a manifestation of the modulatory principles proposed here and may require a different explanation. To support flexible behavior and action planning, sensory computations must go beyond merely representing stimuli with high accuracy and must engage in more complex computations, for example, detection of sensorimotor mismatches (*59*), surprise detection (*60*), or perceptual inference (*61*). Although the current work does not focus on more complex computations, we foresee that the principles proposed here could extend to objectives other than efficient representation of stimuli.

### Outlook

Our results complement previous theoretical studies that identify common themes in sensory systems of diverse organisms, such as the encoding of auditory spatial cues (*62*) or the sampling of natural scenes through saccadic eye movements (*54*). By highlighting the existence of simple principles that can explain different strategies of sensory coding used by various species, these studies offer the hope of developing a general theory of information processing by biological sensory systems.

## METHODS

### Video recordings

We recorded videos of a variety of natural environments while the camera was stationary and moving. Using a GoPro Hero10 Black mounted on an extension pole, we mimicked the perspective of a mouse by holding the camera close to the ground. We recorded ten videos, each with a duration of approximately 1 minute, in five different environments for each locomotion condition for a total of 100 minutes of video data divided evenly between stationary and moving conditions. For the stationary recording, we held the camera in roughly the same position and the field of view was weakly fluctuating. For the moving recordings we walked with the camera in a straight line at an average speed of approximately 50 cm/s. For the purpose of an additional analysis, we recorded videos at different speeds (see below). For all recordings, the camera was set to a linear digital lens with motion stabilization turned off. All videos were recorded in 1080p HD, 1920 by 1080 pixels (px), at 30 frames per second (fps) with a 16:9 aspect ratio. The field of view (FOV) of the camera is approximately 92 (horizontal) by 61 (vertical) degrees (deg) according to online [documentation](https://community.gopro.com/s/article/HERO10-Black-Digital-Lenses-FOV-Informations?language=en_US). Hence, there are approximately 20 px per 1 deg of visual angle. Before recording in a new environment, the exposure of the camera was automatically computed and fixed for all videos in that environment.

### Gabor filters

Before applying filters, each video was normalized by subtracting the mean and dividing by the standard deviation across all pixels over time. To analyze statistics of natural videos we used a large set of parameterized Gabor filters of the following form: 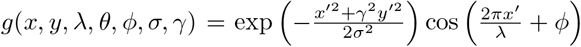, where *x*^′^ = *x* cos *θ* + *y* sin *θ* and *y*^′^ = − *x* sin *θ* + *y* cos *θ*. In this equation, λ denotes the spatial wavelength (inverse of the spatial frequency), *θ* defines the orientation, *ϕ* is the phase, *σ* is the standard deviation of the Gaussian envelope, which controls the spatial extent of the filter, and γ is the spatial aspect ratio. For all filters, we set γ = 1. We considered a range of orientations *θ* ∈ 0, {22.5, 45, 67.5, 90, 112.5, 135, 157.5} deg and phases *ϕ* ∈ {0, 90, 180, 270} deg. The envelope size was determined by the spatial wavelength of the filter i.e. *σ* = λ/3. Filters were replicated at 9 different positions in the scene at 1/4, 1/2, and 3/4 of both the horizontal and vertical dimensions. Depending on the analysis, we considered different numbers of filters with different distributions of spatial frequencies, ranging from 0.02 to 6 cycles per degree (cpd). Details are described in the corresponding sections below.

### Variance of filter outputs

We split the time series signal of each filter output for every video into overlapping excerpts of length *τ* ∈{1, 2, 5, 10, 20 }seconds that were shifted by a single sample. In each of these windows we computed the standard deviation (SD) of the filter outputs. After grouping the data by locomotion condition and spatial frequency, we computed the mean SD over time windows and across various filter orientations, phases, and spatial positions.

### Nonlinearity optimization

We simulated the response of neurons by passing the outputs of the Gabor filters, denoted by s, through a logistic nonlinearity: 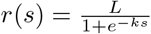, where *L* is the maximal firing rate and *k* is the slope. Assuming that the neuron adjusts its offset to match the mean of the filter output in a short time window, we only consider the second order moment and approximate the distribution of filter outputs using a Gaussian distribution with the same variance. We identified the optimal parameters *k*^∗^ and *L*^∗^ that maximize the utility *U*(*k, L*) = *I*(*s*; *r*) −λ_*r*_ ⟨ *r* ⟩ for a range of Gaussian stimuli, where the standard deviation *σ* was matched to the empirically observed range of SDs from the windowed filter outputs. Here, *I*(*s*; *r*) is the mutual information between the stimulus and the response, ⟨*r*⟩ is the mean firing rate, and λ_*r*_ is a constraint parameter. Specifically, we evaluated 280 *σ* values spaced logarithmically between 10^−1^ and 10^3^, and for each, performed a grid search over 200 logarithmically spaced *k* values from 10^−3^ to 10^−1^ and 200 linearly spaced *L* values from 0.05 to 10. This process was repeated for 20 different constraint strengths λ_*r*_ ranging linearly from 0.5 to 10. Since the mapping from stimulus to response is deterministic, *H(r* | *s*) = 0 and we need only consider the response entropy for the mutual information computation, *I(s; r*) = *H(r*) − *H*(*r* | *s*) = *H*(*r*). Discretizing the stimulus and response values with a fixed binning, we directly computed the probability for each stimulus bin using the Gaussian probability density function. Using the logistic function, the stimulus bins were mapped to response bins and thus the probability for each response bin was computed as a sum of probabilities of the stimulus bins that were mapped to that response bin. We used 10^5^ stimulus bins ranging from − 3 × 10^3^ to 3 × 10 ^3^ and used 10^5^ response bins ranging from 0to 10. The discrete entropy of the response was calculated as 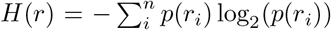, where *r*_*i*_ denotes the midpoint value of the *i*-th bin of the response values. The mean firing rate was calculated directly from the probability distribution of the response bins and the midpoint value of each bin: 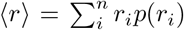 The optimal nonlinearities for each filter were selected by choosing the *k*^∗^ and *L*^∗^ optimized for the standard deviation *σ* of that filter’s output.

### Temporal statistics of filter outputs

To understand how the temporal statistics change during locomotion we computed the Fourier power spectrum of Gabor filter responses. We did so for every filter in each video in short time windows of 5 seconds. Before computing the spectrum in each window we subtracted the mean of that window. The windows overlapped by 2.5 seconds. The spectra were then averaged across time windows and videos for each filter for the stationary and moving conditions separately.

### Temporal filtering

We modeled the temporal filtering performed by a model neuron as a low-pass filter with a linear roll-off in the frequency domain. The filter was thus defined as a piecewise linear function:

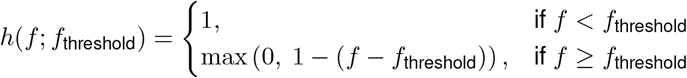

where *f*_threshold_ denotes the frequency threshold beyond which higher frequencies are attenuated. We computed the average spectra of the outputs of individual Gabor filters (stimuli), as well as the spectra averaged across the entire Gabor filterbank and normalized the stationary and moving spectra by the same constant such that the total power of the stationary spectra was equal to one. The spectra of stimuli were then multiplied by the filter *h*(*f*; *f*_threshold_) to generate output spectra. We considered 1000 frequency thresholds ranging linearly from 0 to 15 Hz, which was equal to the Nyquist limit. The total power of the filtered spectra were then computed for each value of *f*_threshold_ . Using the total power of the stimulus in the stationary condition as a reference, we allowed for the model neuron to increase the amount of spectral power transmitted during locomotion: 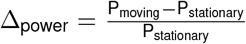, where *P*_moving_ and *P*_stationary_ are the total stimulus power in moving and stationary conditions respectively. We considered 300 different values of Δ_power_ ranging linearly from 0.01 to 3. For each value of Δ_power_ we identified the corresponding frequency threshold *f*_threshold_. The frequency thresholds computed for stationary and moving conditions were denoted *f*_*s*_ and *f*_*m*_ respectively. Using the Wiener-Khinchin theorem, we computed the autocorrelation function of the output of temporal filters in Fig. 3d to a broad-band white noise input.

### Optimization of inhibitory connections

To determine a population of model receptive fields, we performed Independent Component Analysis (ICA) using the FastICA algorithm (*63*) on 32 by 32 px image patches from videos down-sampled by a factor of 8. For each video 5,000 patches were chosen from random positions in random frames, resulting in 500,000 image patches. Each image patch was normalized to have zero mean and unit variance. We learned 101 independent components (ICs). Thus for each reference neuron the neighboring population had 100 model cells. The learned ICs have spatial frequencies ranging from 0.08 to 0.48 cpd (as determined by numerically fitting Gabor filters). The IC filters were then applied to each video in the same manner as the parametric Gabor filters described above. We then randomly selected 1,000 windows of length *τ* = 5 seconds from across all of the videos for each filter. To simulate neuronal activity *r*_*i*_ of each model neuron *i*, the output of each IC filter was demeaned and rectified to enforce positivity. Within each stimulus window, we computed the correlation coefficients ρ_*ij*_ between all pairs of simulated neuron responses *r*_*i*_, *r*_*j*_ in the stationary and moving conditions. In order to remove correlations between filter pairs we considered a simple model of a network interaction where the simulated neural responses *r*_*i*_ of all neurons in the population are weighted and subtracted from a reference neuron *r*_*j*_ to minimize correlation i.e 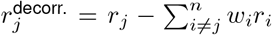, where 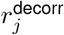 is the response of the neuron r_*j*_ after the decorrelation transform. We optimized the weights 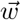 independently for each window and reference neuron such that the average correlation strength was minimized while keeping the weights small: 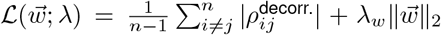 Here 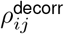 is the Pearson correlation between the post-decorrelation neural response of the reference neuron *r*_*j*_ and population neighbors r_*i*_. λ_*w*_ = 0.1 is a weak regularizer of the weight magnitude. We used a gradient based optimization to find the weights which minimized the loss function.

### Comparisons to published data

The theoretical predictions for the mouse visual cortex use filters which match the range of spatial frequencies of receptive fields in mouse V1 reported in (*32*). Unless stated otherwise, the spatial frequencies of the filters range linearly from 0.02 to 0.32 cpd with an increment of 0.01 cpd resulting in 8,928 filters. For all results the constraint strength λ_*r*_ on the mean responses was set to 1 and the optimal nonlinearity parameters were chosen based on SDs computed from windows with a size of 5 seconds.

### Contrast tuning curves

We simulated the response of optimized model neurons to scaled Gabor patches, identical to the receptive field of the analyzed neuron (i.e. optimal stimuli). The scaling parameter emulated changes in the stimulus contrast. We varied the scaling parameter from 5 to 500 logarithmically, and computed the contrast tuning curves using optimal nonlinearity parameters in stationary and moving conditions. For each filter the contrast tuning curve was normalized by the maximal value between the stationary and moving conditions. The scaling parameter was also normalized by the maximal value and converted to a percentage.

### Orientation tuning curves

To generate orientation tuning curves, each model neuron was exposed to Gabor stimuli identical to the receptive field of the analyzed neuron. Stimuli were presented at 16 different orientations *θ* linearly spanning 0 to 337.5 deg. Each stimulus was also scaled by a fixed contrast parameter to avoid saturating the nonlinearity, while still evoking a response. The filter outputs to the different orientations were then passed through the optimal nonlinearity for each filter to generate tuning curves. The additive and multiplicative gain terms were fit by regressing the moving tuning curve from the stationary tuning curve as in (*8*). A contrast parameter value of 30 was used in the example tuning curve shown in Fig. 2f. For the additive and multiplicative gain terms a contrast parameter value of 10 was used. We considered nonlinearities optimized for 5,472 Gabor filters with spatial frequencies ranging linearly from 0.02 to 0.02 cpd.

### Dependence of locomotion modulation on movement speed

Videos were recorded in three different environments at three different average speeds (20 cm/s, 30 cm/s, and 50 cm/s). We recorded two 60 second videos for each speed and environment resulting in the total of 360 seconds per speed. We computed the output of the Gabor filterbank, and the optimal nonlinearities of model neurons were found using the method described previously. The optimal nonlinearity parameters for the stationary videos were used for the zero speed condition. The response of optimal nonlinearities at each speed were computed to 10^4^ samples of Laplace-distributed stimuli with µ = 0 and a scale of *b* = 10. The response for each filter was then averaged across stimulus presentations. The mean response for each model neurons as a function of movement speed was then fit using an exponential function r(v) = *a* −*be*^*cv*^. To limit the computational cost we only used filters with frequencies ranging from 0.01 to 0.12 resulting in 3,456 filters.

### Population coding fidelity

To simulate neural responses, we used nonlinearities optimized for a population of low spatial frequency Gabor filters of various orientations, phases, and positions approximately matching the distribution of spatial frequencies reported in (*13*) (spatial wavelength ranged from 0.01 to 0.12 resulting in 3,456 filters). We computed the response of this population to 2 × 10^5^ samples of stimuli sampled from the Laplace distribution with a mean of *µ* = 0 and a scale of *b* = 50, using nonlinearities optimized for the moving and stationary states. We then split the data in half and trained a linear decoder on the first half of the data to predict the stimuli from the population response vector. Using the held out data, we split the stimuli into non-overlapping blocks of 1,000 samples and in each block computed the average response of the population and root mean squared error of the decoder. To simulate a switch from stationary to locomotion the first half of the stimulus blocks simulated for the stationary condition were concatenated with the second half of the stimulus blocks for the moving condition. The resulting trajectories of average firing rate and decoding error were then smoothed with a Gaussian filter with *σ* = 1 windows.

### Temporal dynamics

The data from (*6*) were reanalyzed by computing the autocorrelation of the normalized response of individual mouse V1 neurons to moving dot stimuli in stationary and locomotion conditions. The autocorrelations were computed on the average response of each neuron to each stimulus speed thus generating the number of response profiles equal to the number of neurons times the number of stimulus speeds. For the theoretical results, we allowed a 50% increase in power during locomotion compared the total power during the stationary condition, and extracted the optimal *f*_threshold_ for this constraint. Applying the Wiener-Khinchin theorem we then computed the autocorrelation by taking the inverse Fourier transform of the power spectra of temporal filters responding to white noise stimuli, as described above. For each analyzed autocorrelation function, we computed the decay time, defined as the lag time where the autocorrelation decreases to or below 0.5. We performed a paired t-test on the decay times of individual neurons in the stationary and locomotion conditions to determine if there was a significant change in the temporal processing during movement. The same analysis was performed for simulated neurons.

### Surround suppression

We first computed the response of each independent model neuron (i.e. an IC filter followed by rectification) to stimuli of increasing size. The stimuli were full-field gratings with the orientation, phase and spatial frequency matched to the IC filters by fitting parametric Gabor filters. We presented grating stimuli of different sizes, determined by the radius of a circular mask. We tested 40 radii ranging form 0 to 16 deg. To simulate the surround suppression we used the weights optimized to minimize correlation magnitude in the stationary or moving conditions and for each model neuron we subtracted the weighted response of the population for the stimuli of increasing size. To compare to individual neurons we used weight vectors 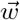 optimized to example stimulus windows. To compare to the population average we averaged across all stimulus windows. We computed a suppression index in the same manner as in (*14*). Finally, we computed the population response as a function of stimulus size by first normalizing the response of each reference filter by the maximal value across both locomotion conditions and then averaging across filters.

### Gain modulation in mice and marmosets

To perform an interspecies comparison we relied on distributions of spatial frequencies of receptive fields in mouse V1 and foveal receptive fields in the marmoset V1 reported in (*32*). For each of these distributions, we generated a population of 2,000 neurons whose corresponding Gabor filters had spatial frequencies sampled accordingly. The orientation, phase, and position of each filter were independently sampled from a uniform distribution with limits described above. For each of these Gabor filters we used the pre-computed optimal nonlinearity parameters for the stationary and moving conditions. We simulated neural responses by exposing model nonlinearities to 1,000 Laplace-distributed stimuli with *µ* = 0 and *b* = 50. The same process was repeated for the comparison to peripheral V1 neurons in the marmoset where the distribution of spatial frequencies in the far periphery was based on the data from (*33*).

We quantified the change in the shared, multiplicative gain in populations of model neurons at the transition to locomotion in the following way. We first simulated responses to the identical set of random stimuli described above using nonlinearities optimized for locomotion and the stationary state. This resulted in two matrices of simulated neural activity *X*^*m*^ and *X*^*s*^ where an entry 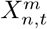 or 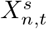 corresponded to the response of the *n*-th neuron to the *t*-th stimulus during movement or in the stationary state, respectively. To compute the change in shared multiplicative gain in both states, we performed linear regression of the form *X*^*m*^ = *a*^*m*^*X*^*s*^, where *a*^*m*^ is a scalar multiplicative factor that affects the activity of all neurons when the behavioral state changes. By definition, the value of the multiplicative factor in the stationary state is equal to 1 i.e. *a*^*s*^ = 1. We performed this analysis for populations of nonlinearities optimized to mouse-like, low-frequency and primate-fovea-like, high-frequency Gabor filters. In both cases the linear fit was reasonable with the coefficient of the goodness of fit *R*^2^ equal to 0.794 and 0.997 respectively, which implies that this simple model indeed emulates well a shared gain change in the population of simulated neurons. We report differences in of the gain terms *a*^*m*^ and *a*^*s*^ = 1 in Fig. 5e.

### Analysis of data from freely moving mice

We reanalyzed the publicly accessible data from (*15*) to determine how locomotion affects the stimulus statistics experienced by freely moving mice. Using videos from the head-mounted “world-view” camera, which records the visual scene from the animal’s point of view, we applied a Gabor filter bank with parameters identical to those used in the analysis of the natural videos (see above). Since the video had a different resolution (40 by 30 px) and spanned a larger FOV of about 120° a different scaling factor of 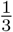 px per degree of visual angle was used. The running speed data is occasionally missing values, so we only considered windows with complete data. The average running speed in each 5 second window was then computed and the data were split based on the first quartile of the average running speed into stationary (< 0.42 cm/s) and moving (> 0.42 cm/s) conditions. The SD of filter outputs in each window was then computed and for each locomotion condition we evaluated the average SD. The Fourier transform of the filter outputs was determined for each window and we then took the average over all windows and filters. We then considered correlations between the responses of independent component filters in 5 second windows. Due to the low resolution of the video data, we performed ICA again on 16 by 16 px image patches from the natural videos down-sampled by a factor of 20 using a similar procedure as above. Since the dimensionality was initially reduced with PCA to retain 75% of the variance, only 48 ICs were learned. The learned IC filters have spatial frequencies ranging from approximately 0.01 to 0.33 cpd (as determined by numerically fitting Gabor filters). As before, the output of each IC filter was demeaned and rectified to emulate neural responses. The correlation between all pairs of IC responses in each window was then computed and the average correlation magnitude was computed for each locomotion condition.

### Minimal model of sensing during locomotion

We constructed a minimal model to explain the changes in signal statistics during locomotion. We modeled a point-agent that observes stimuli while moving in a spatially heterogeneous two-dimensional environment. The environments were created by convolving a two-dimensional Gaussian filter with i.i.d. white noise in a 100 by 100 grid. The spatial extent of the Gaussian filter *σ*_spatial_ determines how spatially correlated or smooth the resulting environment is. The output of the convolution was then rescaled to have a maximal value of 100 and a minimal value of -100. At each time step of the simulation, the agent observed a stimulus which was normally distributed with the mean specified by the grid position occupied by the agents sensor (see below) and with the standard deviation *σ* = 0.1. The position of the agent 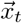 was governed by the following equation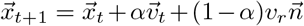, where *α* is a momentum term, *v*_*t*_ is the current velocity of the agent, and *v*_*r*_ is the translational speed that controls the variance of the stochastic 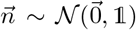 sampled from a two-dimensional multivariate normal distribution (*64*). We set *α* = 0.5. The starting position *x*_0_ was randomly drawn from a uniform distribution over grid positions, and the initial velocity *v*_0_ was assigned unit norm with a direction drawn uniformly over all angles. We simulated different sensors (corresponding to e.g. overlapping receptive fields), by allowing the agent to sample from 8 spatial positions equally distributed in a circle of radius *r* from its current position. For each combination of the speed and spatial scale, we simulated ten different trajectories of length 1,500 time steps for ten different environments. For the simulated results in Figures 1-4, *σ*_spatial_ = 5 was used and for the stationary condition *v*_*r*_ = 0.2 in comparison to the moving condition where v_*r*_ = 5. For Figure 5 we considered spatial scales of *σ*_spatial_ = 2 and *σ*_spatial_ = 0.5 and adjusted the moving velocity to *v*_*r*_ = 1. For all simulated results we used a radius of *r* = 10 for the sensors. For the inset in Fig. 2f, we used five speed values *v*_*r*_ ∈ {0.10.2, 1, 2, 5}.

## ACKNOWLEDGEMENTS

We thank Laura Busse, Ann Hermundstad, Steffen Katzner and Ruben Portugues for helpful discussions and suggestions.

## Funding

The authors acknowledge that they received no funding in support of this research.

## Author contributions

Conceptualization: J.M.G. and W.F.M. Methodology: J.M.G. and W.F.M. Software: J.M.G. Validation: J.M.G. and W.F.M. Formal analysis: J.M.G. and W.F.M. ınvestigation: J.M.G. and W.F.M. Resources: W.F.M. Data curation: J.M.G. and W.F.M. Writing - original draft: J.M.G. and W.F.M. Writing - review and editing: J.M.G. and W.F.M. Visualization: J.M.G. and W.F.M. Supervision: W.F.M. Project administration: W.F.M. Funding acquisition: W.F.M.

## Competing interests

The authors declare that they have no competing interests.

## Data, code, & materials availability statement

All data and code needed to evaluate and reproduce the results in the paper are freely available. The data (natural videos) used in the analysis are available at https://doi.org/10.12751/g-node.14m4gq. The code is publicly available at https://doi.org/10.5281/zenodo.20624372. No new materials were generated in this study.

## SUPPLEMENTARY MATERIALS

**Figure S1:**
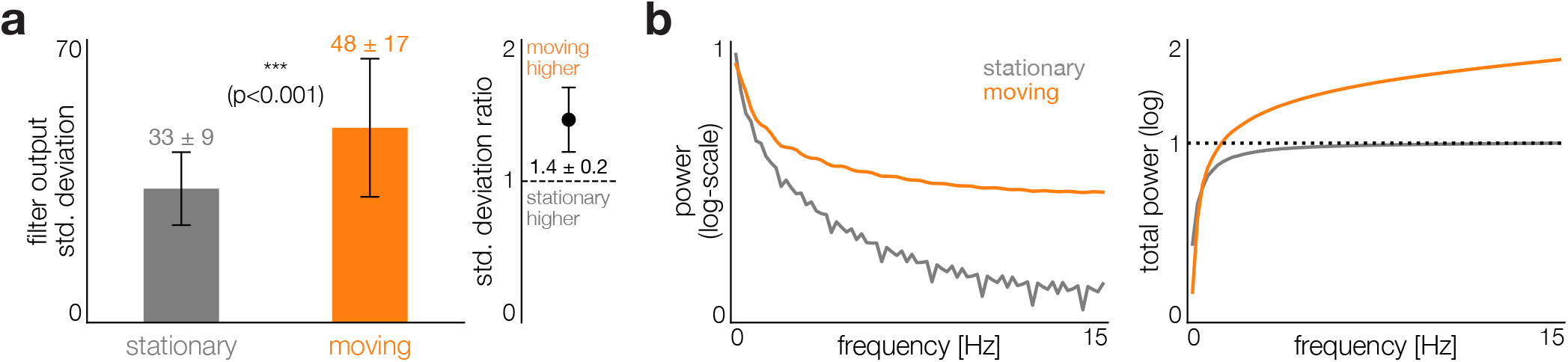
Simulated eye movements do not qualitatively change the impact of locomotion on sensory statistics. We simulated random eye movements that match the distribution of saccade sizes and frequencies observed in mice (*54*). Specifically, the center point of the filter was modified at fixed time intervals (1 second for moving videos and 2 seconds for stationary videos) by adding a randomly signed variable sampled from a Gaussian distribution with a mean of 10 degrees and standard deviation of 3 degrees in the horizontal dimensions and a mean of 1 degree and standard deviation of 0.3 degrees in the vertical dimension. The positions of the filters were clipped such that the filter lay within the bounds of the video frame. **a)** Left panel: Average standard deviation of a large Gabor filterbank where the spatial frequency ranged from 0.03 to 0.12 cpd in stationary (gray) and moving (orange) conditions. The numerical value of the average and the standard deviation of the average are above the bars. The average standard deviation in the moving condition was significantly higher than in the stationary condition (paired t-test, t-statistic=81.27). Right panel: ratio of the average standard deviation of Gabor filter outputs during movement to the average during the stationary condition. The numerical value of the average ratio and the standard deviation of the average ratio are below the point. Error bars indicate standard deviation. **b)** Left panel: the average power spectrum of the outputs of a large Gabor filterbank where the spatial frequency ranged from 0.03 to 0.12 cpd during locomotion (orange) and stationary (gray) conditions. Right panel: the total power (i.e. the integral) of the average spectrum as a function of the frequency for the stationary (gray) and moving conditions (orange).

**Figure S2:**
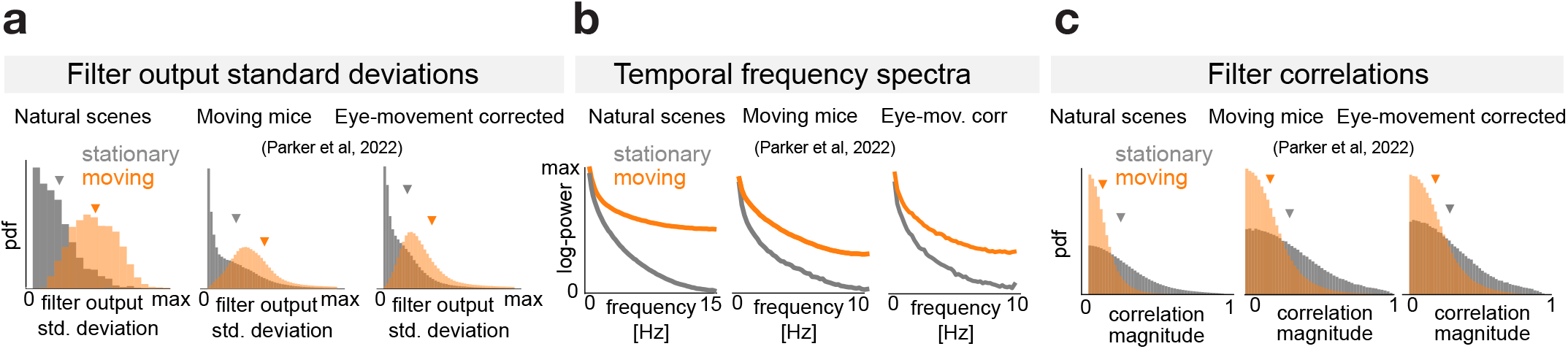
Comparison of distributions of statistical features between natural scene recordings and videos from the perspective of moving mice before and after correction for eye movements (data from (*15*)). To correct for the eye position, for each mouse we trained the “shifter network” that maximizes the predictive strength of a Generalized Linear Model (GLM) of the neural population activity. We used the code and followed the procedure described in (*15*). A range of L2 regularization parameters for the GLM weights (*λ* ∈[0.01, 1000]) were used and the network with the lowest test loss was used to correct the “world-view” camera videos for eye movements. Using the eye-movement corrected videos, we followed the same procedure as in the analysis of data from freely moving mice (see above). Correcting for eye movements in freely moving mice does not qualitatively change the impact of locomotion on the statistics of sensory inputs. **a)** Standard deviations of filters (triangles denote averages). **b)** Temporal spectra of filter outputs. **c)** Magnitudes of pairwise correlations between filter outputs (triangles denote averages).

**Figure S3:**
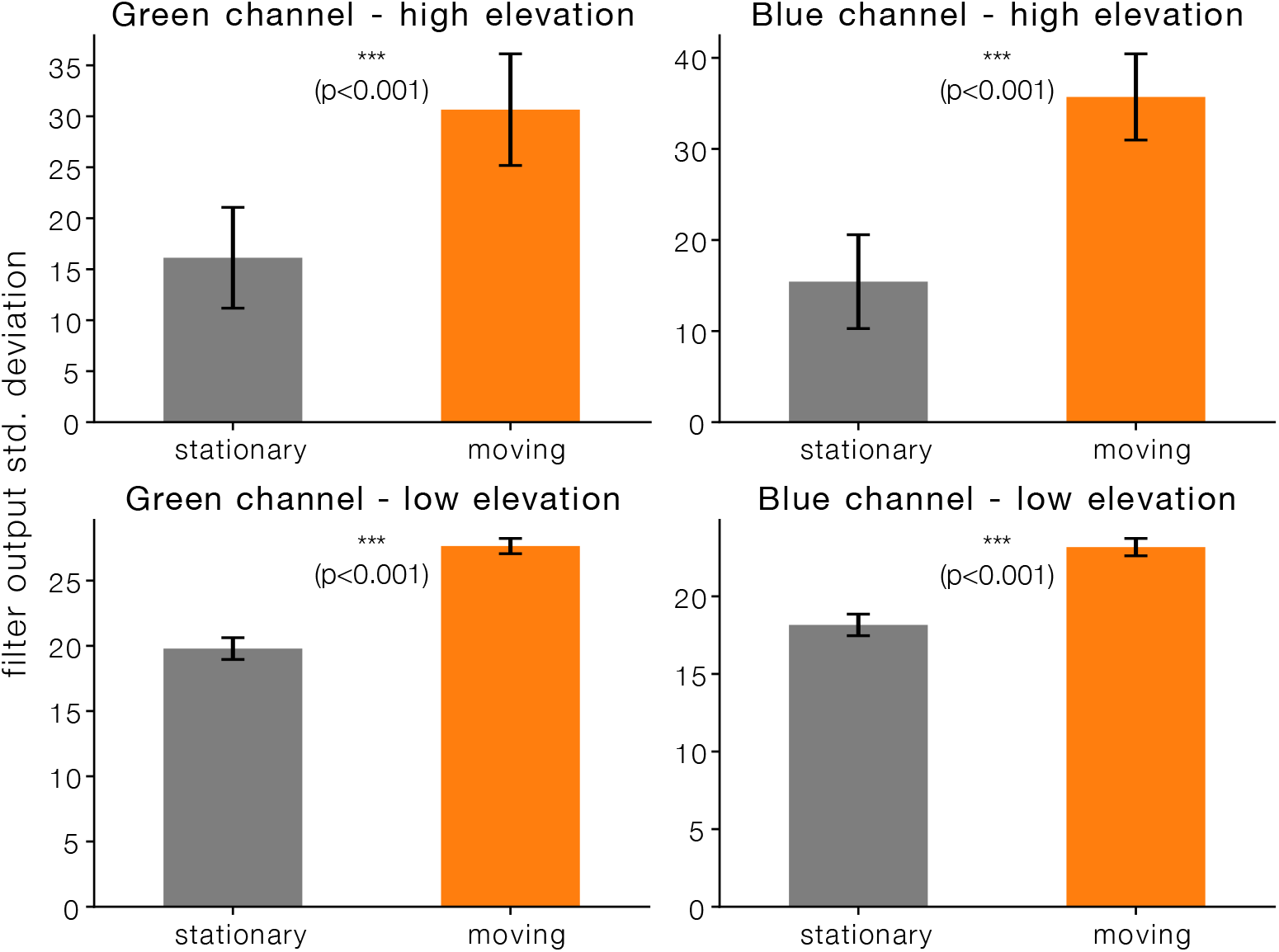
Color vision and elevation do not qualitatively change the impact of locomotion on the standard deviation of filter outputs. The green and blue channels of the natural videos were extracted and the outputs of a bank of Gabor filters were computed for videos recorded in 5 different natural environments in two movement conditions (for a total of 10 videos). The filterbank spanned 8 orientations *θ* ∈{0, 22.5, 45, 67.5, 90, 112.5, 135, 157.5} deg, had a fixed phase of zero, and a spatial frequency of 0.1 cpd. The outputs were computed from two positions in the visual scene: high and low elevation (1/4 and 3/4 of the vertical dimension respectively, centered in the horizontal dimension). Following the same analysis explained in the methods, the filter outputs were segmented into 5 second windows and the standard deviations per window computed. The average standard deviation of filter outputs was computed across windows and environments for each movement condition, color channel, orientation and position and are summarized by the bar plots which are organized by color channel and position and report the mean of the average standard deviations across filter orientations. The columns show the results for the green channel (left) and blue channel (right). The rows show the results for the high elevation (top) and low elevation (bottom). For all color channels and elevations, the filter output standard deviations are significantly higher while moving than while stationary (paired t-test, t-statistic: 33.854 (upper left), 53.256 (upper right), 19.006 (lower left), 17.113 (lower right)).

**Figure S4:**
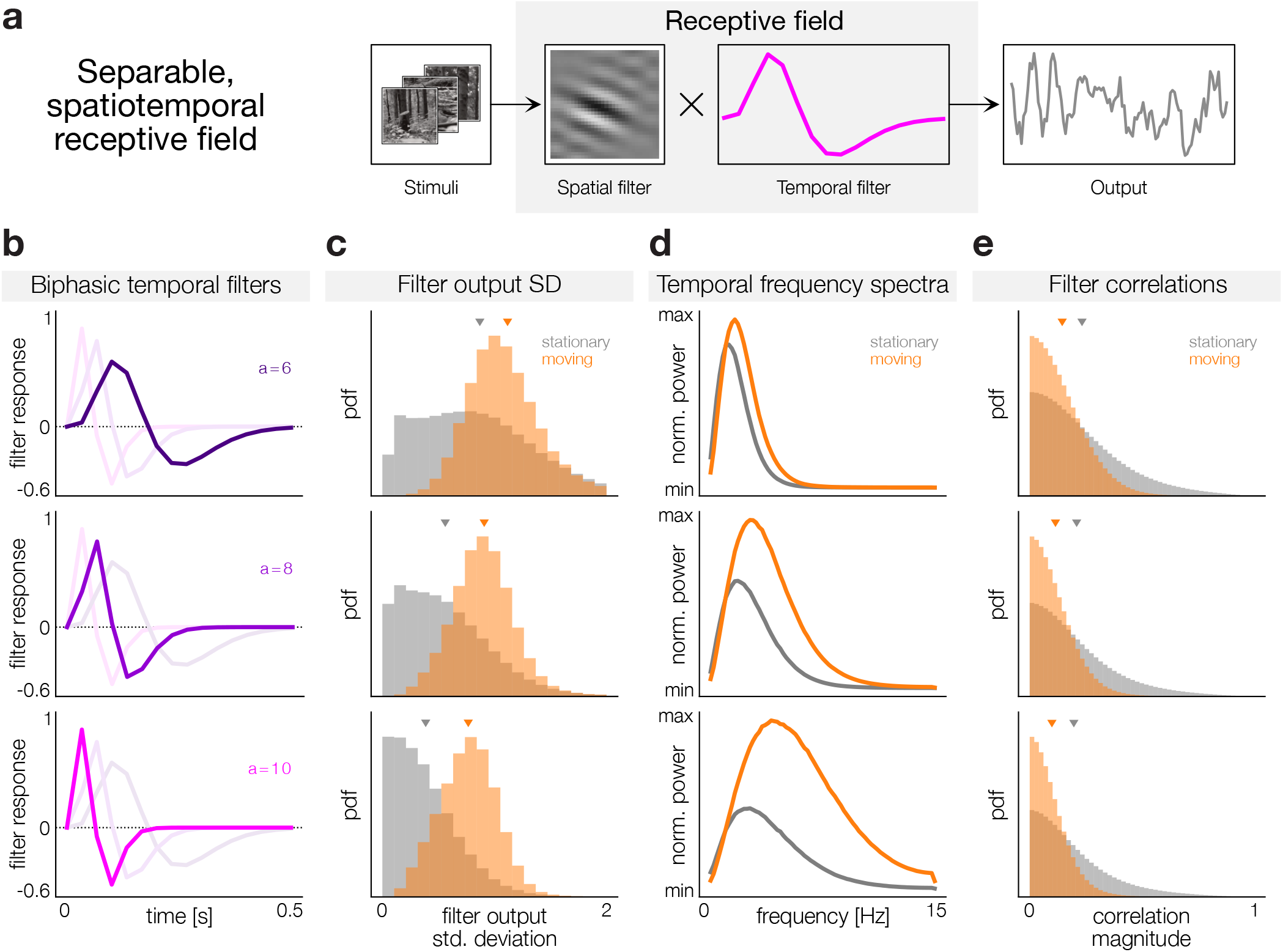
Outputs of spatiotemporal filters are systematically influenced by of locomotion. The outputs of the spatial ICA filters were temporally filtered with biphasic temporal filters reminiscent of temporal receptive fields. The biphasic temporal filters are parameterized as 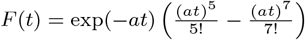, where the parameter *a* controls the timescale of the filter as in (*65*). We used temporal filters with a length of 0.5 seconds and three values of *a* ∈{6, 8, 10} to match the timescales of temporal receptive fields in mice. The temporally filtered outputs were then processed in the same manner described earlier for the analysis of the Gabor filterbank outputs and ICA filters to compute the filter output standard deviations, temporal spectra, and filter correlations. **a)** Schematic showing the application of a separable, spatiotemporal receptive field to video stimuli. **b)** Biphasic temporal filters with three different timescales. **c)** Standard deviations of temporally filtered ICA filter outputs (triangles denote averages). For all values of *a* the average filter output standard deviation is higher during locomotion than in the stationary condition consistent with the previous results (Fig. 2). **d)** Temporal spectra of temporally filtered ICA filter outputs. **e)** Magnitudes of pairwise correlations between temporally filtered ICA filter outputs (triangles denote averages).

**Figure S5:**
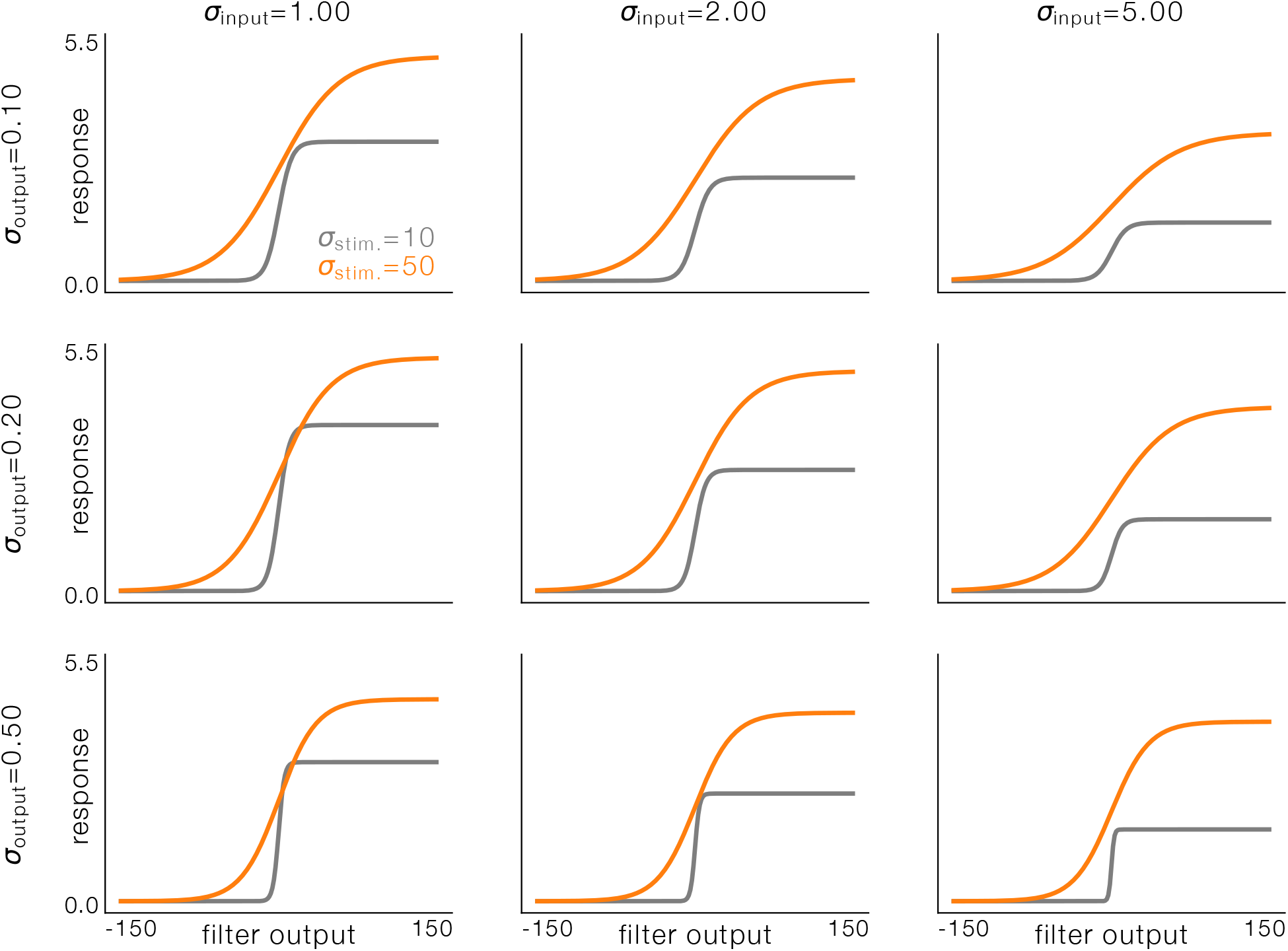
Noise does not qualitatively impact the gain increase of optimal nonlinearities during locomotion. Optimal nonlinearities were computed for low and high variance stimuli for a range of input and output noise levels. For both the input and output noise Gaussian random variables were used. The stimulus standard deviations of *σ*_stim._ ∈ *{*10, 50*}* were chosen to roughly approximate typical values of filter output standard deviations to stationary and moving videos respectively. The standard deviations of the input noise used were *σ*_input_ ∈ *{*1, 2, 5*}* and the output noise standard deviations used were *σ*_output_ ∈{0.05, 0.1, 0.2 }. For the input signal, 10^6^ samples were randomly chosen from a Gaussian distribution. A grid search was performed over the nonlinearity parameters with 100 logarithmically spaced *k* values from 0.01 to 1 and 100 linearly spaced *L* values from 0.1 to 10. The mutual information was estimated directly from the discretized joint and marginal distributions of the stimuli and firing rates with a fixed binning scheme with 2, 000 bins ranging from −150 to 150 for the stimuli and 2, 000 bins ranging from 0 to 10 for the firing rates. The average firing rate and utility were computed in the same manner as before with *λ*_*r*_ = 0.5. With increasing input noise the maximal firing rate of the nonlinearities tends to decrease so as not to amplify the input noise. With increasing output noise, the slope of the nonlinearities tends to increase, that is to say they get steeper.

**Figure S6:**
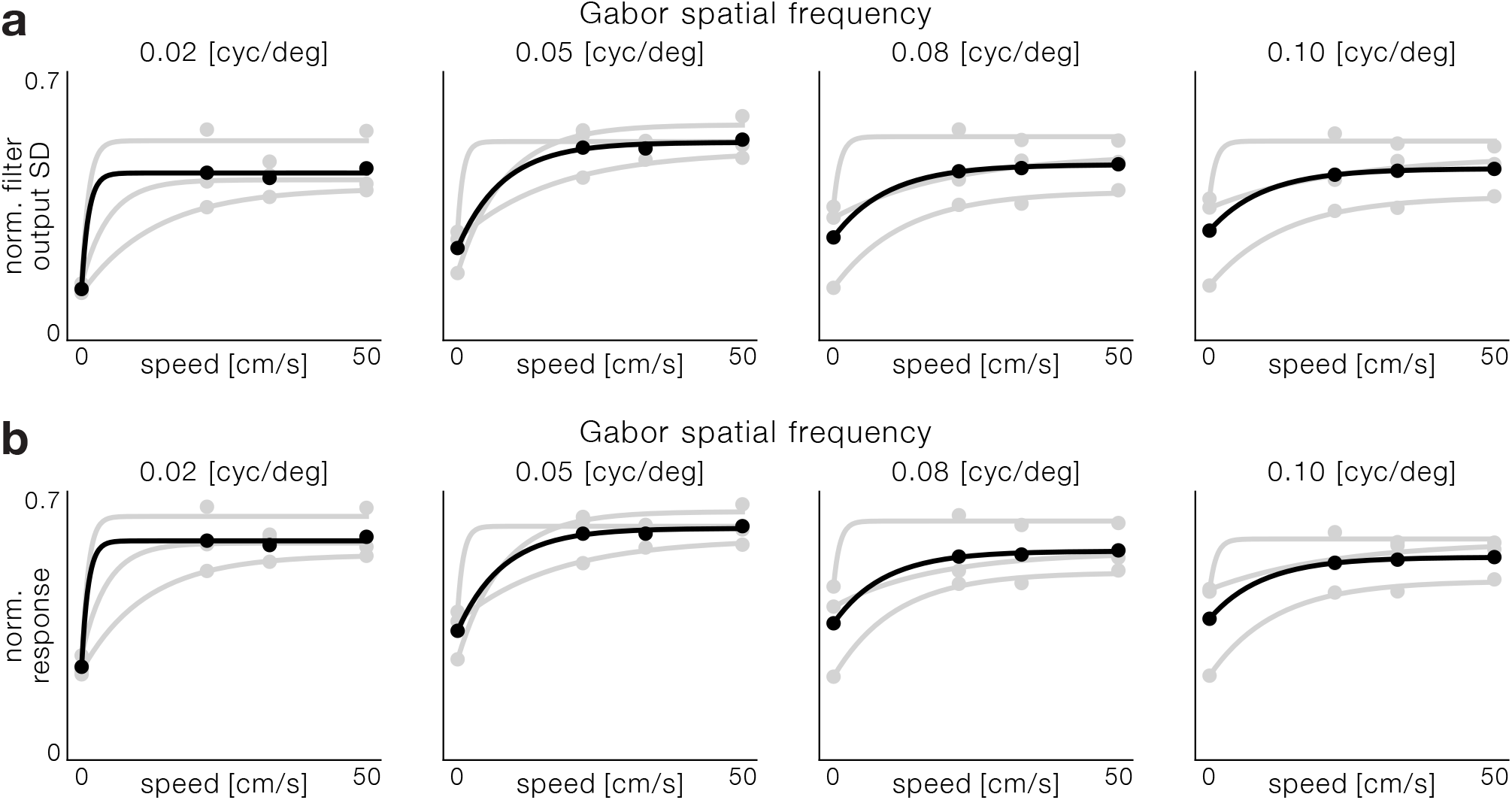
Locomotion increases firing rates at the onset of locomotion across different spatial frequencies and environments. The same procedure as in the methods section was followed for videos taken at 3 different speeds in three different environments: forest, field, and orchard. Rather than taking the average of the standard deviation of filter outputs across environments, the average standard deviation of filter outputs was computed for each environment separately. The average standard deviation as a function of movement speed was then fit with an exponential function as before for each environment. For comparison the mean over the average standard deviations per environment and speed were also computed and fit with an exponential function. The optimal nonlinearities were chosen and the firing rates were computed in a similar manner as before, but per environment rather than over all environments. Lastly the mean over the average firing rates per environment was computed and fit with an exponential as well. **a)** Average standard deviations of a bank of Gabor filters as a function of running speed for different spatial frequencies. Gray lines/points are for individual environments and the black line/points are averaged over environments. **b)** Average normalized response of the optimal nonlinearities for a bank of Gabor filters as a function of running speed for different spatial frequencies. Gray lines/points are for individual environments and the black line/points are averaged over environments. For this analysis *λ*_*r*_ = 1.

## Notes

### Competing Interest Statement

The authors have declared no competing interest.

### Summary of Updates

This is the accepted version of the manuscript

